# Intracranial brain stimulation modulates fMRI-based network switching

**DOI:** 10.1101/2021.01.12.426446

**Authors:** Mangor Pedersen, Andrew Zalesky

**Author notes:** **Corresponding author:** School of Clinical Sciences, Auckland University of Technology (AUT), Private Bag 92006, Victoria St West, New Zealand.

## Abstract

The extent to which functional MRI (fMRI) reflects direct neuronal changes remains unknown. Using 160 simultaneous electrical stimulation (es-fMRI) and intracranial brain stimulation recordings acquired in 26 individuals with epilepsy (with varying electrode locations), we tested whether brain networks dynamically change during intracranial brain stimulation, aiming to establish whether switching between brain networks is reduced after intracranial brain stimulation. As the brain spontaneously switches between a repertoire of intrinsic functional network configurations and the rate of switching is typically increased in brain disorders, we hypothesised that intracranial stimulation would reduce the brain’s switching rate, thus potentially normalising aberrant brain network dynamics. To test this hypothesis, we quantified the rate that brain regions changed networks over time in response to brain stimulation, using *network switching* applied to multilayer modularity analysis of time-resolved es-fMRI connectivity. Network switching and synchrony was decreased after the first brain stimulation followed by a more consistent pattern of network switching over time. This change was commonly observed in cortical networks and adjacent to the electrode targets. Our results suggest that neuronal perturbation is likely to modulate large-scale brain networks, and multilayer network modelling may be used to inform the clinical efficacy of brain stimulation in epilepsy.

## Introduction

Network modularity encompasses a family of algorithms that quantify whether a collection of network nodes exert stronger ‘intramodular’ connectivity than expected by chance – i.e., modularity provides a set of subnetworks with stronger than average within-network connectivity (Sporns and Betzel, 2016). Multilayer network modularity represents a multidimensional version of network modularity (De Domenico, 2017; Mucha et al., 2010). A multilayer network is conceptualised as a ‘network of networks that are connected across several dimensions (Bassett et al., 2013; Betzel and Bassett, 2017; Vaiana and Muldoon, 2018). This is an explicit modelling framework that allows information to be shared across edges connected, for example in space and time, enabling us to track *where* and *when* entities in a network transit between different sub-networks or modules. In turn, spatiotemporal network measures like multilayer modularity are promising approaches that can enhance our understanding of human brain function, and a way to monitor the clinical response to invasive treatment strategies, including intracranial brain stimulation.

By extending the concept of modularity to several dimensions, es-fMRI connectivity can be represented in terms of a spatiotemporal network model of the brain (Finc et al., 2020; Lydon-Staley et al., 2018; Shine et al., 2016). Multilayer network flexibility, or switching, is associated with cognitive functions including working memory (Braun et al., 2015), reasoning (Pedersen et al., 2018a), reward (Gerraty et al., 2018) and fatigue (Betzel et al., 2017) as well as alterations in multiple psychological and neurological disorders (Gifford et al., 2020; Harlalka et al., 2019; Long et al., 2019; Paban et al., 2019; Shao et al., 2019; Tian et al., 2020a). There is also evidence that brain network switching changes in response to behavioural training. For example, Bassett et al. (2011) showed that motor training is associated with greater network switching, particularly in association cortices involved in higher-order cognition. A follow-up study demonstrated that brain network switching on the first day of motor training was correlated with individual differences in overall motor learning rate (Telesford et al., 2017). Another study showed increased brain network switching in people who underwent half a year of musical training, compared to people with no musical training (Li et al., 2019). These studies suggest cause-and-effect relationships such as overt learning (e.g., motor, and musical training) can alter the brain’s network switching rate.

It remains unknown whether focal neuronal perturbation –for example via invasive brain stimulation– leads to observable changes in the rate at which the brain switches between es-fMRI networks. In this study, we used the above-mentioned multilayer modularity model (Mucha et al., 2010) to investigate spatiotemporal network changes that occur during short periods of intermittent brain stimulation. Such spatiotemporal network models are advantageous for this purpose as they can pinpoint the specific time points when brain regions transit between networks. Elucidating network switching during brain stimulation will provide new insights into the dynamical properties of es-fMRI networks.

To test whether brain network switching is altered during intracranial brain stimulation, we used es-fMRI data with concurrent intracranial stimulation acquired from individuals with focal epilepsy. Focal epilepsy is a neurological disease associated with seizures arising from a circumscribed part of the brain that in some cases progress from a focal to bilateral tonic-clonic seizure (Fisher et al., 2017). Focal epilepsy is associated with increased fMRI connectivity (Bernhardt et al., 2011, 2015; Hong et al., 2017; Pedersen et al., 2015, 2016, 2017), and prior electrophysiology research in epilepsy suggests that the impact of brain stimulation results in changes to neuronal networks and provides clinical benefits (Gummadavelli et al., 2015; Khan et al., 2009; Schulze-Bonhage, 2017; Toprani and Durand, 2013; Zangiabadi et al., 2019) likely by ‘steering’ the brain into a temporary state associated with modulated network events (Li and Yang, 2017). High-frequency stimulation (here, 100 Hz) is thought to be associated with neuronal inhibition (see Garcia et al. 2005, for a review) and neuronal inhibition is also associated with attenuated fMRI activity (Aksenov et al., 2019). We believe once the fMRI activity of brain regions is attenuated due to high-frequency stimulation, it is less likely that the brain regions can switch between different modular networks or states. Consequently, we hypothesise that es-fMRI network switching decreases after brain stimulation, in people with focal epilepsy.

## Materials and Methods

### Participants

We studied 26 patients with treatment-resistant epilepsy for whom 160 es-fMRI scans were acquired while simultaneously receiving 100Hz intracranial electrical stimulation (Figure A.1 – see also Oya et al., 2017 and Thompson et al., 2020). Intracranial brain stimulation is a common part of the pre-surgical work-up in treatment-resistant focal epilepsy patients. The most common stimulation target was the amygdala followed by the Heschl’s gyrus and the frontal/cingulate gyri. For the most part, participants were implanted with multiple electrodes, as shown in Figure 1. All es-fMRI datasets were downloaded from openneuro.org (https://openneuro.org/datasets/ds002799/versions/1.0.3). The dataset is described in detail in Oya et al. (2017) and Thompson et al. (2020). Open-access MATLAB-based scripts for computing instantaneous phase synchrony, multilayer modularity and network switching can be found at https://github.com/MangorPedersen/fMRI_codes/.

**Figure 1:**
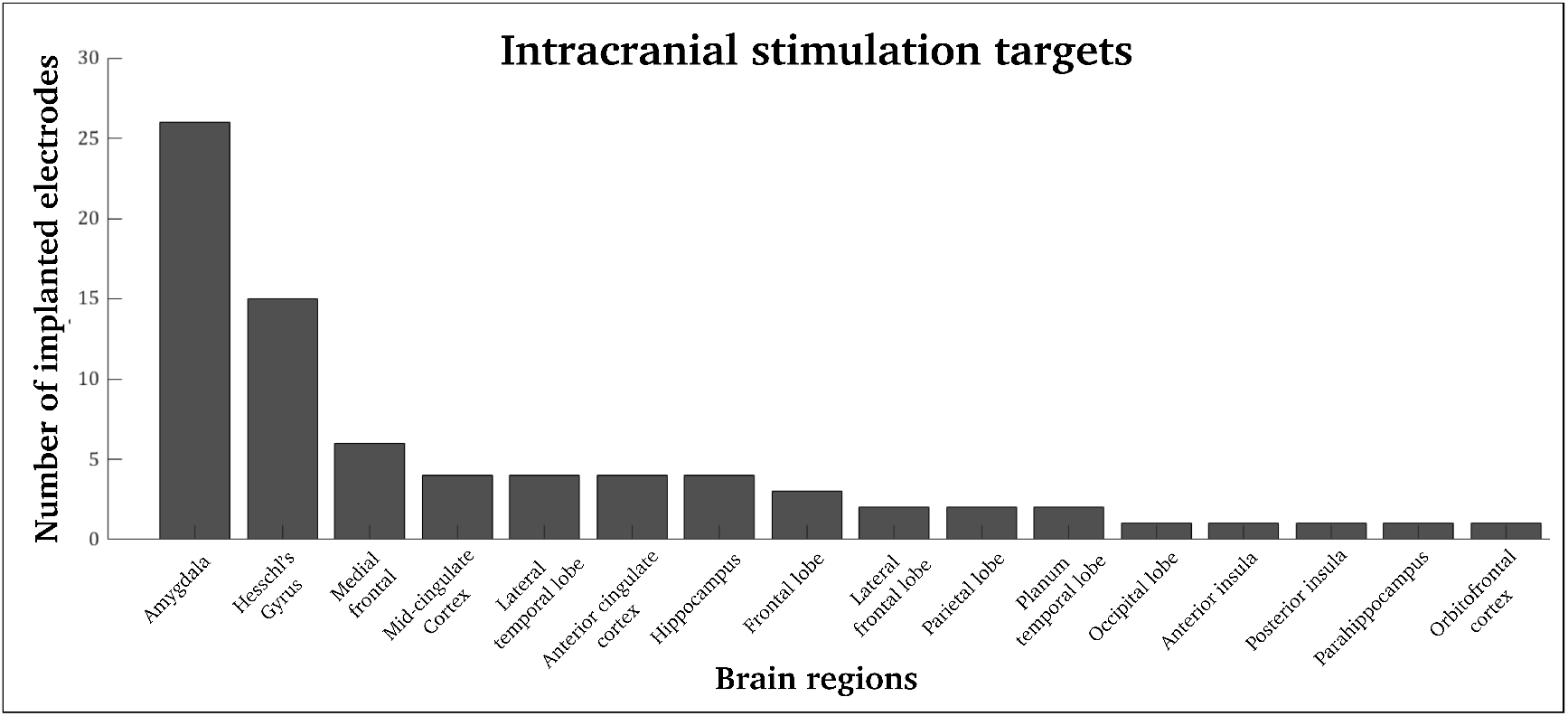
Location of brain electrodes across 26 epilepsy patients: The most common stimulation site included the amygdala (26 electrodes) followed by Heschl’s gyrus (15 electrodes) and the frontal/cingulate cortex. Note that most individuals were implanted with more than one electrode.

### es-fMRI and brain stimulation parameters

The length of each es-fMRI scan varied between participants. We included es-fMRI scans that were more than 9 minutes long (160 scans in total). The es-fMRI echo time was 30 milliseconds, and the data had a voxel size of 3×3×3 millimetre. There was no significant difference in head motion between NoStim and Stim epochs (Thompson et al., 2020). The repetition time of the es-fMRI data was 3000 milliseconds with a delay in repetition time of 100 milliseconds. The electrical stimulation was delivered during this repetition time delay and ensured no artefacts between MRI radiofrequency coils and electrodes (Oya et al., 2017). The brain stimulation consisted of bi-phasic charge-balanced square pulses (50-90 milliseconds in length, 8-12 milliamps, and 5-9 pulses at a 100 Hz stimulation rate). Pre-processing performed using fMRIPrep – see Appendix A, for full details about fMRI preprocessing in this cohort.

### Filtering and parcellation of es-fMRI data

The pre-processed es-fMRI data was zero-phase filtered within narrow-band frequencies of 0.03 and 0.07 Hz, using a 5^th^ order Butterworth filter (‘*filtfilt’* function in MATLAB). After filtering the es-fMRI in the forward direction, we also filtered the data in the reverse order to minimise distortion and filter effects at the beginning and end of the signals (Dwivedi and Vyas, 2011). Narrow-band filtering is a requirement of instantaneous phase synchrony, to satisfy the Bedrossian’s theorem when using the Hilbert transform for time-series analyses (Honari et al., 2020).

The average fMRI signal within 196 brain regions was extracted for analysis. We combined the cortical parcellation mask from the Human Connectome Project (Glasser et al., 2016) with 180 bilateral cortical regions and the sub-cortical parcellation mask from Tian et al. (2020b) with 16 bilateral sub-cortical regions (Figure A.2). We used a symmetric brain parcellation mask –i.e., the same parcel in homologous brain regions– to counter the laterality effects of temporal lobe epilepsy where seizures predominantly originate from a single hemisphere (Adcock et al., 2003). For each es-fMRI scan, this results in a 3D tensor of size 196 x 196 x 160 that represents the interconnectivity between each of the 196 brain regions, for 160 es-fMRI time-points (80 Stim time-points and 80 NoStim time-points, partitioned into alternating epochs – see Figure 2). In line with Thompson et al. (2021), we excluded the first three time-points from each epoch to avoid overlap between Stim and NoStim, resulting in a total of 112 es-fMRI time-points for analysis.

**Figure 2:**
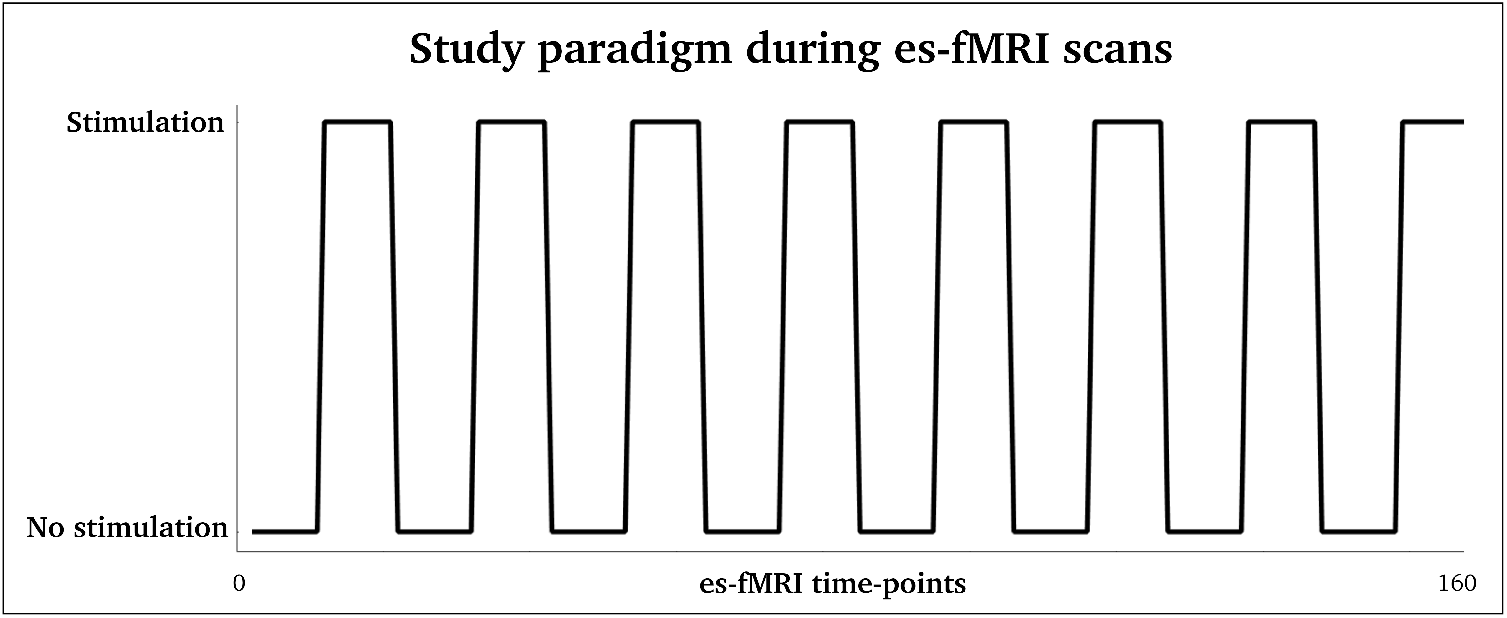
Study paradigm used during es-fMRI scans: es-fMRI scans had 10 data-points with NoStim (30 seconds) followed by 10 data points of Stim (30 seconds), repeated eight times (16 epochs in total). This resulted in 80 Stim data points (4.5 minutes) and 80 NoStim data points (4.5 minutes), for each es-fMRI scan. After excluding the first three time-points from each epoch, we analysed 112 time-points.

### Instantaneous Phase Synchrony

As shown in Figure 2, an es-fMRI block design was used in this study with alternating 30 seconds NoStim epochs followed by 30-second Stim epochs. To quantify time-varying fMRI connectivity within relatively short half-a-minute epochs, we used instantaneous phase synchrony. Instantaneous phase synchrony quantifies narrow-band fMRI connectivity by estimating the phase difference between brain regions, at a single time-point resolution (Glerean et al., 2012; Pedersen et al., 2018b; Ponce-Alvarez et al., 2015).

Instantaneous phase synchrony is calculated by using the Hilbert transform (Bedrosian, 1963) to extract phase information between all brain regions (see Figure A.3). In the equation below, *Y* is a 2D matrix comprising the average narrow-band es-fMRI data across 196 brain regions and 160 time-points, using the same procedure as reported in Glerean et al. (2012); Pedersen et al. (2018b); and Ponce-Alvarez et al. (2015). 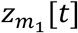 and 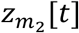 is the analytic representations of the rows in *Y* (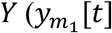 and 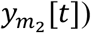):

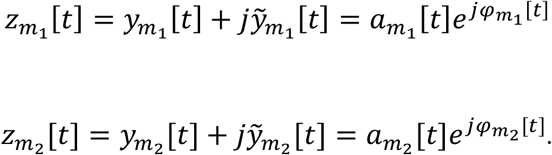

Here, 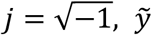 represents the Hilbert transformation of *y* where 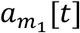 ad 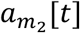 is the instantaneous amplitudes. 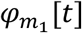 and 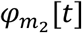 being the instantaneous phases of 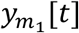 and 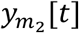, respectively. 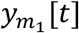 and 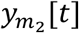 are phase-locked if:

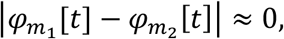

with | . | as an absolute value operator. The instantaneous phase difference between 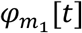 and 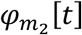 is derived from the rows in the analytic matrix *Z*.

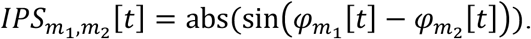

The angle information between signals allows us to quantify their phase synchrony by estimating the absolute sinusoid difference between the signals, at each time point, *t*. We then computed 1 minus phase synchrony (1 – *IPS*) to obtain a numerical range of phase coherence between 0 and 1. A value of 0 indicates no phase synchrony between two brain regions, and a value of 1 indicates that two brain regions are fully synchronous (Mormann et al., 2000).

It remains debated what sparsity level matrices contain optimal levels of biological information and a minimal influence of noise confounders (Fornito et al., 2012; van den Heuvel et al., 2017; Langer et al., 2013). To minimise potential confounders in this study, we thresholded and binarized matrices at several density thresholds, retaining 10%, 15%, and 20% of the strongest instantaneous phase synchrony connection pairs. In Figure A.4, we provide the minimum phase synchrony values (i.e., threshold cut-off values) for each of these thresholds.

### Multilayer modularity and network switching rate

To quantify the rate of brain network switching we first generated a multilayer modularity model (Mucha et al., 2010). The multilayer modularity model is based on the Louvain modularity algorithm (Blondel et al., 2008; Lancichinetti and Fortunato, 2012):

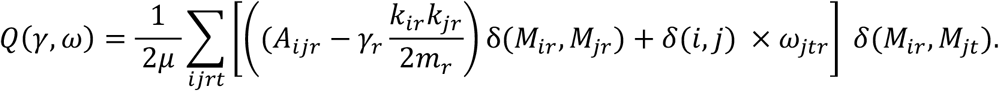

The input to the multilayer modularity model is *A* and denote the thresholded and binarized 3D instantaneous phase synchrony tensors, for each es-fMRI scan. *A*_*ijr*_ is the phase synchronisation between brain region, *i* and *j*, for time-point *r*. *k* is the degree (the total number of connections) at brain region *i* at time-point *r*, and *m* refers to the total degree across the 196 brain regions at timepoint *r*. The Newman-Girvan null model of intra-network connectivity (2*m*_*r*_) is used to quantify whether the intramodular degree is greater than expected by chance (Sarzynska et al., 2016). Topological modularity is controlled by the network resolution parameter, *γ*_*r*_, at time-point *r*. We used *γ* = 1, 1.1, 1.2 and 1.3 in this study where low *γ* values will return larger and fewer brain modules whereas high *γ* values will return smaller and more brain modules. Temporal connectivity is controlled by the coupling parameter, *ω*_*jtr*_, at brain region *j* between adjacent layers (i.e., adjacent time-points) *r* and *t*. We used *ω* temporal coupling parameters of 0.1, 1, 2 and 3 (in line with recommendations from Yang et al., 2020).

δ(*M*_*ir*_, *M*_*jr*_) and δ(*M*_*ir*_, *M*_*jt*_) has a value of 1 if two brain regions of interest (*i*, *j*) are located within the same module, and 0 if they are allocated to two separate modules (Bassett et al., 2013). Modularity maximisation methods are inherently heuristic (Good et al., 2010), and our multilayer modularity models converged after an average of 4 iterations across all es-fMRI scans. The modular decomposition across all participants is displayed in Figure A.5 and the most common modules that resembled i) the visual network, ii) the somatomotor network, iii) the frontoparietal network, iv) the default-mode network and v) sub-cortical brain regions. The average network modularity was calculated with the *Q*-value, which range between 0 connections (no modular structure and between-module connectivity) to 1 (fully modular structure and within-module connectivity). Q-values and number of modules for all network densities (10%, 15% and 20%) as well as *γ* (*γ* = 1, 1.1, 1.2, and 1.3) and *ω* (*ω* = 0.1, 1, 2, 3) values are found in Figure A.6.

We used instantaneous phase synchrony as an input to an ordinal multilayer modularity network model, enabling us to quantify the percentage of times each brain region changes network allegiance –*i.e., network switching*– during intracranial brain stimulation. The network switching rate is derived from the multilayer modularity model and can be written as:

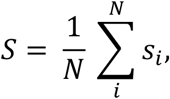

where *s*_*i*_ is the network switching at node *i*, calculated as the number of times a brain region transits between network modules, divided by the total number of possible network transitions (Bassett et al., 2011). *S* was calculated within each *NoStim* and *Stim* epoch, excluding the first three time-points to avoid overlap between epochs.

### Analysis

All findings are based on thresholded/binarized instantaneous phase synchrony tensors at a 15% network density (i.e., retaining the top-15% synchronous connection-pairs, and a topological/temporal modularity resolution of *γ*/*ω* = 1, consistent with previous studies (Bassett et al., 2013). Replication analyses for multiple network densities (van den Heuvel et al., 2017; Langer et al., 2013) and multiple recommended *γ*/*ω* parameters (Yang et al., 2020) are also reported.

As reported in Figure 3, we used one-way within-subjects (repeated measures) ANOVA to infer differences between all epochs. Post-hoc comparisons between epochs were computed with Bonferroni correction with a critical probability value of *p* < 0.05. We used a one-tail univariate paired-samples t-test to infer the reduction in network switching for the 196 brain regions between the first epoch (no brain stimulation) and the average of all remaining epochs. We used false discovery rate correction at *q* < 0.05 (Benjamini and Hochberg, 1995) to control for multiple comparisons across all brain regions (Figure 4).

**Figure 3:**
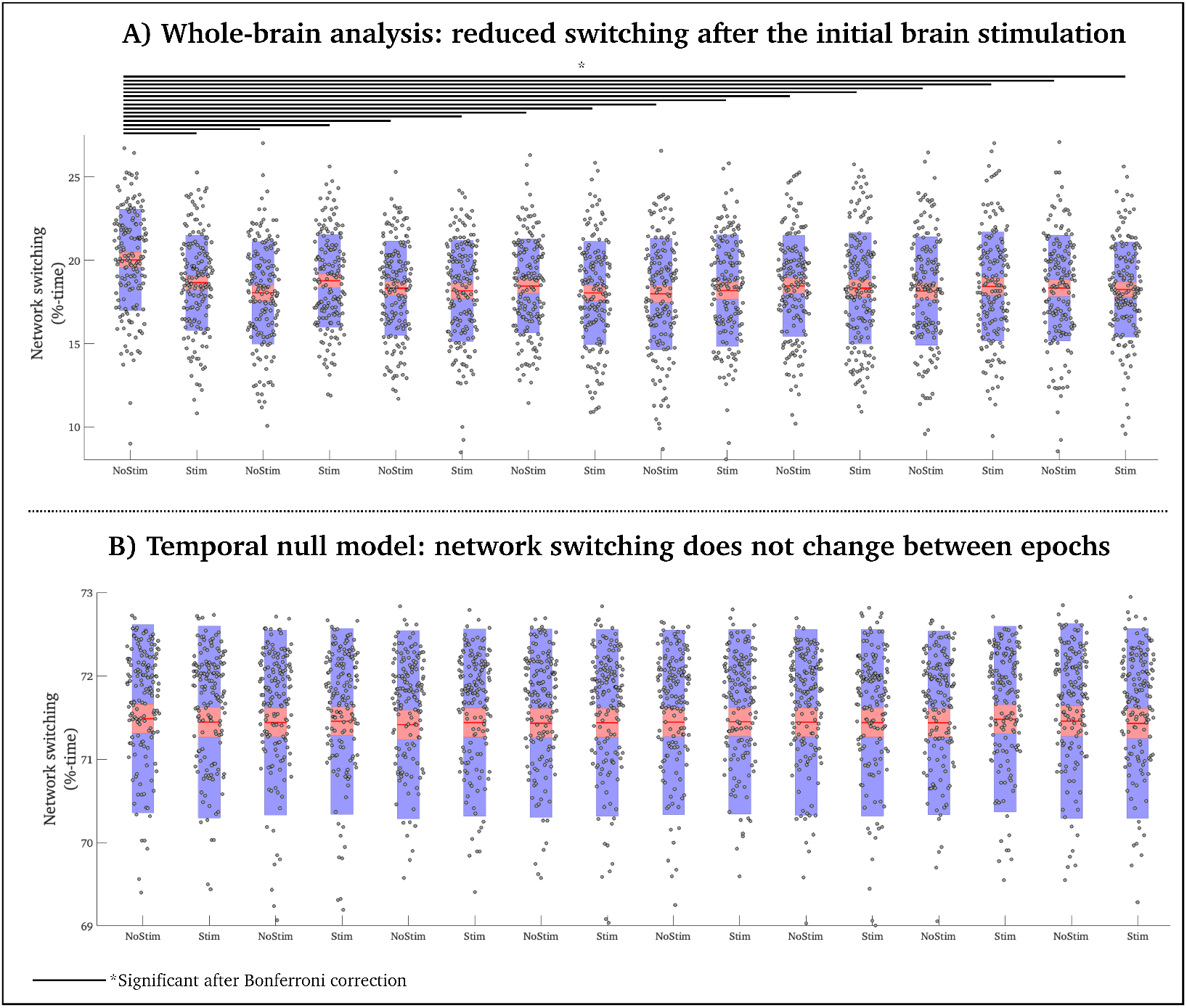
Stimulation-induced changes in network switching at group level: A) Network switching shown as a function of time throughout the es-fMRI scans, stratified according to NoStim and Stim epochs (epoch numbers in parentheses). Here, network switching is the proportion of time that nodes switch network allegiance during Stim and NoStim epochs. The shaded red area is the 95^th^ confidence interval of the mean. The shaded blue area is one standard deviation of the mean. B) An overview of post-hoc comparisons between all epochs. The white numbers inside the bars refer to the epoch numbers that are statistically significant for a reduction in network switching within the current epoch (Bonferroni corrected). C) Same as A, but here the es-fMRI time-points are randomly permuted 100 times before computing the multilayer modularity model – i.e., a temporal null model. We report the mean of the 100 permutations.

**Figure 4:**
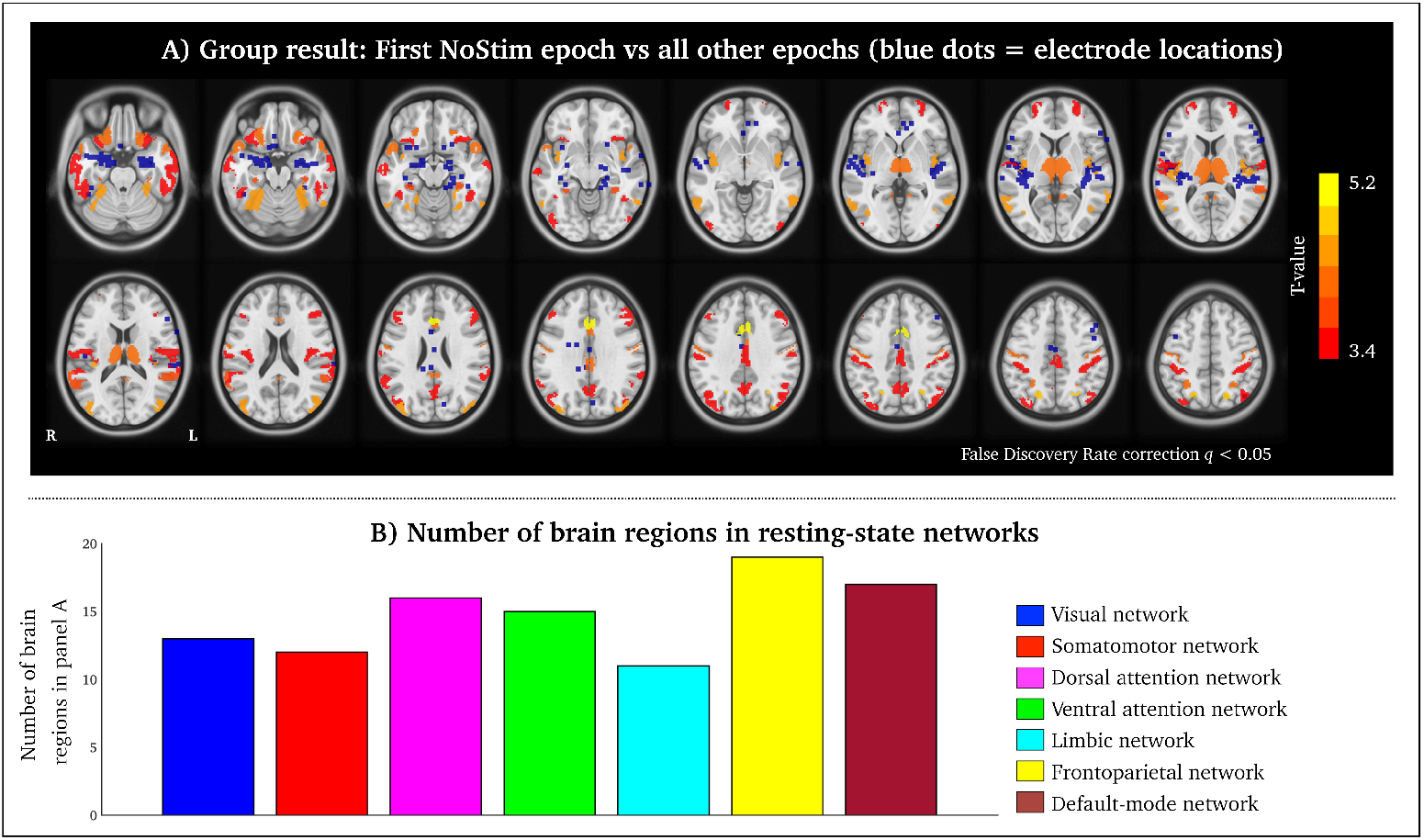
Regional changes in network switching after brain stimulation: A) Brain regions with a significant decrease in network switching after the first stimulation (paired t-test between first NoStim epoch and the average of all other epochs, FDR corrected at q<0.05). Electrodes are highlighted with blue dots. 3/26 individuals in this study did not have coordinates for the intracranial electrodes. B) The number of brain regions from A within seven resting-state networks from Yeo et al. (2011).

We also used Pearson’s correlation coefficient to determine brain regions that positively correlated with the study block design values (Stim/NoStim – see Figure 2). We permuted all network switching and block design values (NoStim time points = 0; Stim time points = 1), 500 times to generate a null distribution. Brain regions that displayed greater correlation than the 95^th^ percentile of the null distribution are displayed in Figure 5.

**Figure 5:**
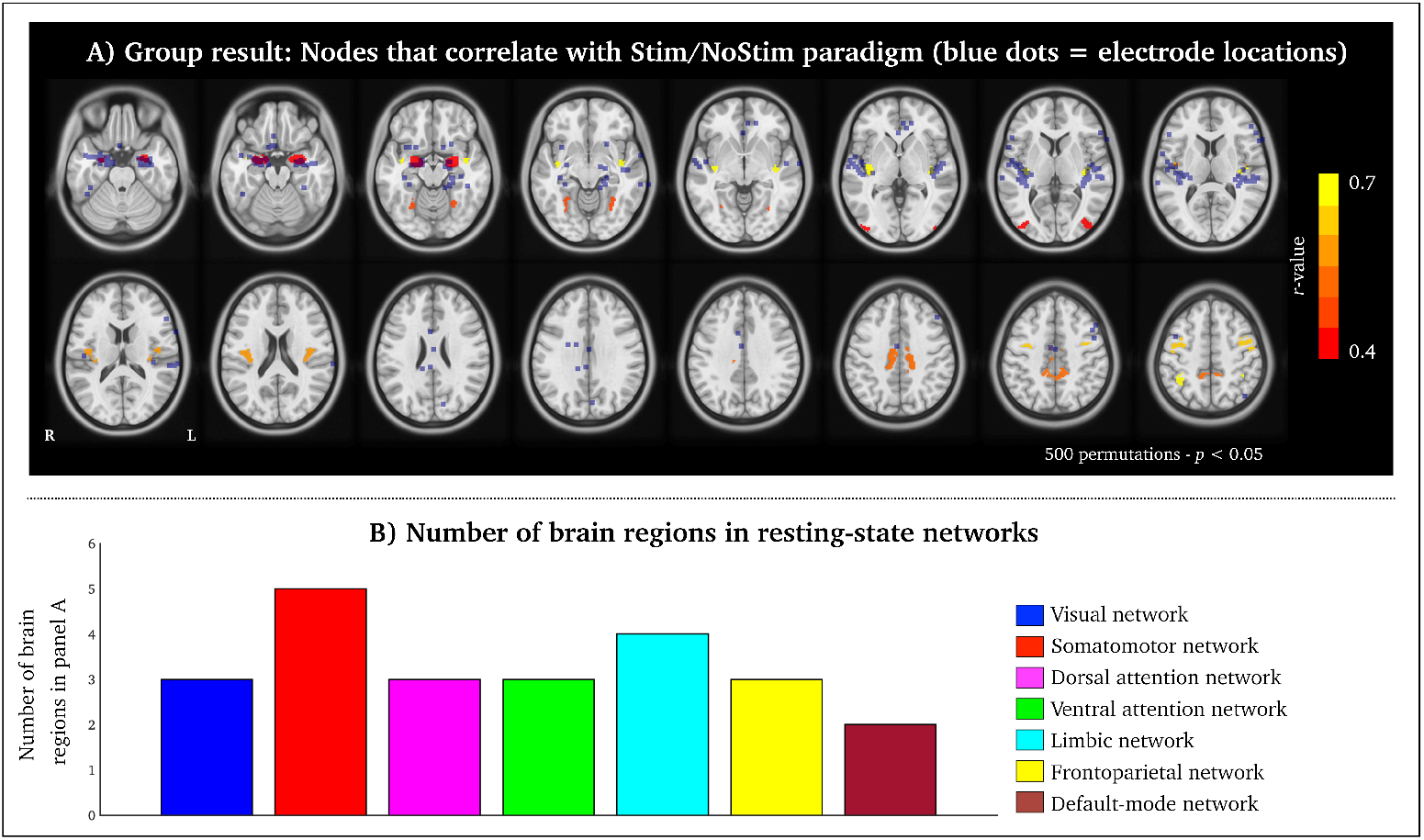
Nodes that correlate with the Stim/NoStim paradigm: A) Brain regions with a greater correlation than the 95^th^ percentile of the null distribution (based on 500 permutations). Electrodes are highlighted with blue dots. B) The number of brain regions from A within seven resting-state networks from Yeo et al. (2011).

To infer individual-specific changes after brain stimulation, we calculated the difference between network switching in the first epoch (NoStim) and the average of all remaining epochs, divided by the standard deviation of all remaining epochs, for each subject with electrode location information available (23/26 epilepsy participants had electrode locations). This resulted in node-specific z-scores for each subject, representing brain regions with stronger network switching in the first epoch compared to subsequent epochs. We averaged network switching from all es-fMRI scans that subjects underwent were averaged before calculating the z-scores (minimum = 2 scans; maximum = 10 scans – see Figure A.1, for full information). We set a z-score threshold of 2.32 corresponding to the 99^th^ percentile of the standard z-distribution.

## Results

### Group-level network switching is decreased after the initial brain stimulation

A one-way repeated measures ANOVA demonstrated a significant main effect of network switching between epochs, *F*(15,2385) = 4.07, *p* < 0.001. Post-hoc tests showed that all the first epoch had greater network switching than all the 15 subsequent epochs (*p* < 0.05, Bonferroni correction – see Figure 3A, for all statistical comparisons), meaning that the main reduction in whole-brain averaged network switching occurred after the first stimulation epoch (Figure 3A – left). 101/160 (63%) of es-fMRI scans had a reduction in network switching after the first stimulation.

We employed a temporal null model previously described by Bassett et al. (2013) to test whether the stimulation-induced changes in network switching could be explained by random dynamical processes. In this temporal null model, we randomly permuted es-fMRI time-points 100 times per es-fMRI scan before computing 100 ‘temporally random’ multilayer modularity models per es-fMRI scan. This procedure preserves the topological modularity of networks but changes their temporal order. As seen in Figure 3B, we observed no differences between epochs when randomly permuting es-fMRI time-points (repeated measures ANOVA (*F*(15,2385) = 0.1, *p* = 1). The network switching was approximately six times greater in the temporal null model (Figure 3B), compared to the original data (Figure 3A). This finding suggests that temporal fluctuations of network switching in response to brain stimulation are unlikely to occur by chance.

### Specific brain regions display decreased network switching during brain stimulation

After establishing that brain-averaged network switching is reduced during brain stimulation, we aimed to delineate brain regions where this reduction was most prominent. We conducted a paired t-test for each of the 196 brain regions, between the first NoStim epoch and the average of all subsequent epochs. Following false discovery rate correction for the 196 tests (Benjamini and Hochberg, 1995), the frontal cortex, parietal cortex, and temporal cortex displayed the strongest reduction in network switching during brain stimulation (Figure 4A). Notably, significant differences in network switching between NoStim and Stim epochs were localized to regions distant from the most common stimulation sites, particularly the amygdala.

Statistically significant brain regions were commonly observed adjacent to, but not overlapping, with intracranial electrodes, suggesting that brain stimulation may influence brain networks beyond the focal stimulation targets (Figure 4 – blue dots). The two brain networks with the most significant brain regions between Stim and NoStim were the default-mode and frontoparietal network (Figure 4B), both spatially distant from the most common stimulation sites. These two networks harbour several integral brain hubs, often referred to as the brain’s rich-club, with strong inter-region connectivity (van den Heuvel and Sporns, 2011). This provides further evidence that focal brain stimulation induces widespread network effects.

Next, we conducted a Pearson’s correlation analysis for each of the 196 brain regions, testing which brain regions positively correlate with the brain stimulation paradigm seen in Figure 2. This correlation analysis quantifies brain regions that display a pattern of decreased network switching during NoStim and increased network switching during Stim. After generating a null distribution using 500 permuted values, we found that the amygdala, parahippocampus, Heschl’s gyrus, cingulate gyrus and frontal cortex positively correlated with brain stimulations. Contrary to the nodal decreases of network switching after the first brain stimulation (Figure 4), the brain regions that correlated with the stimulation paradigm overlapped more with intracranial electrodes, particularly the amygdala and Heschl’s gyrus. This is supported by the resting-state network result showing that limbic and somatomotor networks contained the majority of significant regions (Figure 5).

### Individual-level network switching is decreased during brain stimulation

We conducted an exploratory individual-level analysis by computing a z-score by measuring the standardised difference between network switching in the first epoch (before stimulation) and all subsequent epochs. We observed that all subjects with electrode information available (23/26 subjects) displayed different spatial patterns of network switching after the initial Stim epoch. Notably, all subjects displayed increased network switching proximate, or overlapping, to the individual electrode locations (see blue dots in Figure 6 and Figure A.7).

**Figure 6:**
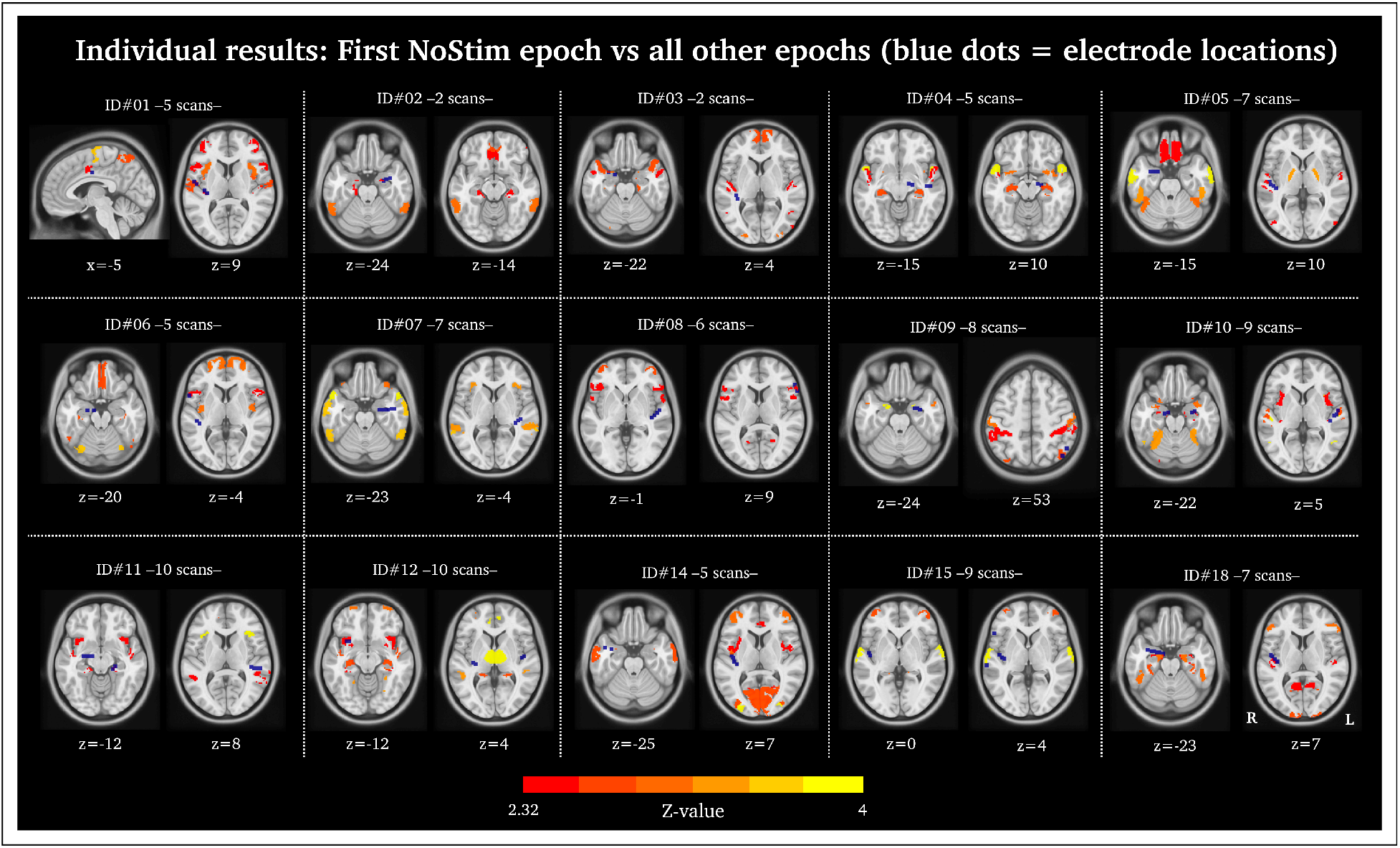
Individual results – first NoStim epoch versus all other epochs: Here we have presented individual-level decreases in network switching occurring after the first NoStim epoch. Included are the first 15 of the 23 epilepsy subjects (from a total of 26 epilepsy subjects) with electrode information available. Electrodes = blue dots. The remaining 8 subjects are presented in Figure A.7.

### Replication across multiple network parameters and densities

We replicated our result of reduced network switching during Stim versus NoStim epochs (FDR corrected) for 16 *γ*/*ω* parameter combinations (*γ* = 1, 1.1, 1.2 and 1.3 and *ω* = 0.1, 1, 2 and 3). We also replicated our results across several proportionally thresholded network densities, preserving 10%, 15% and 20% of phase synchrony connection pairs. Although all network densities were statistically significant between Stim and NoStim epochs (FDR corrected), there was a trend towards greater statistical power between Stim and NoStim epochs at a higher network density threshold with more network connections (Figure A.8).

### Instantaneous phase synchrony is also decreased during brain stimulation

Lastly, we investigated whether the underlying instantaneous phase synchrony between brain regions –i.e., the input data to our multilayer modularity model– was also modulated by the effect of brain stimulation. We found that the mean of instantaneous phase synchrony averaged across all brain regions was reduced during the first NoStim epoch compared to the remaining epochs (paired t-test: *t(159)* = 10.02, *p* < 0.001), using a network density of 15%. Reduced instantaneous phase synchrony may therefore explain the attenuated network switching during brain stimulation (Figure A.9).

In Figure 7 we present three-time series, each representing the instantaneous phase synchrony between two brain regions, where one brain region is located proximate to a stimulation site and the other brain region is distant from a stimulation site. These time series show a negative correlation between brain stimulation and instantaneous phase synchrony and highlights that distant network properties are associated with stimulation targets.

**Figure 7:**
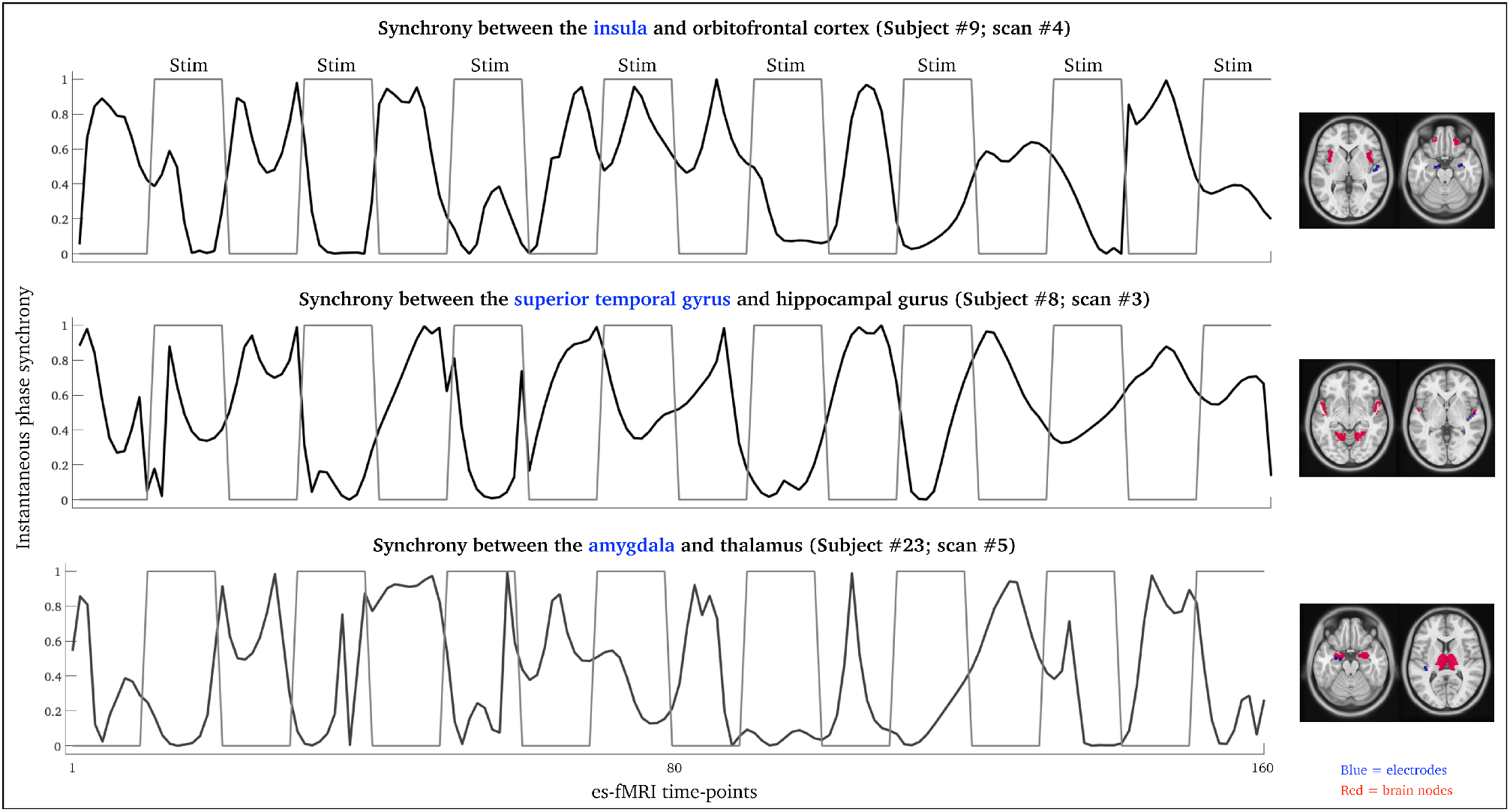
Examples of instantaneous phase synchrony time series: These three time series each represent the synchrony between two brain regions for representative individuals. The blue colour highlight brain regions proximate to an intracranial electrode. The Pearson’s r correlation between the instantaneous phase synchrony time-series and the block design (Stim/NoStim) was −0.62 (p<0.001 – first row), −0.59 (p<0.001 – the second row) and −0.61 (p<0.001 – the third row).

## Discussion

We observed a reduction in brain network switching and synchrony after the initial intracranial brain stimulation (Figure 3A and Figure A.9), which was stable across a range of network parameters (Figure A.8). The initial stimulation may ‘startle’ the brain as the epilepsy patients in this study did not receive stimulation less than 1.5 hours (and up to 3 hours) before the es-fMRI acquisition. In subsequent epochs, network switching stabilized. This implies a lasting network effect beyond the initial impact of brain stimulation and may reflect a long-lasting inhibition of local neuronal activity (Lafreniere-Roula et al., 2010). The initial change in network switching is supported by previous deep brain stimulation Parkinson’s disease showing that tremors are reduced within seconds after administration of brain stimulation to the subthalamic nuclei (Blahak et al., 2009), and the neural effects from a single train of high-frequency stimulation can also last for several minutes (Benazzouz et al., 1995). This suggests that the brain likely reacts rapidly to brain stimulation, and can be maintained for minutes. From a clinical standpoint, our results suggest that stimulation-induced reduction of network switching may normalize aberrant brain dynamics associated with disorders such as epilepsy (Pedersen et al, 2017).

Although the neuronal mechanisms of intracranial brain stimulation are complex, it is well established that invasive brain stimulation affects the cellular, electrical, molecular and network architecture of the brain (Jakobs et al., 2019). Network effects of brain stimulation (i.e., brain changes that occur distant to the stimulation target) are thought to be mediated by large and myelinated axons becoming depolarised and transformed into action potentials (Johnson et al., 2008). In line with our results, Alhourani et al. (2015) suggest that the underlying network mechanisms of intracranial brain stimulation represent a reduction of synchrony between remote brain regions that is achieved by perturbing afferent and efferent neurons that are (directly or indirectly) connected to the stimulation target. This is supported by Middlebrooks et al. (2018) who showed that epilepsy individuals who had a positive clinical response to brain stimulation to the anterior thalamus displayed greater connectivity in the default-mode network, compared to patients who did not respond positively to brain stimulation. This reinforces that brain stimulation impacts widespread brain networks beyond the stimulation target.

Invasive brain stimulation has Class I evidence for seizure reduction in treatment-resistant epilepsy (Li and Cook, 2018), from randomised controlled trials with two different stimulation targets (Fisher et al., 2010; Morrell and RNS System in Epilepsy Study Group, 2011). Brain networks may normalise during invasive brain stimulation and attenuated network switching may represent a putative marker of therapeutic normalisation. This fits the clinical pattern of epilepsy, given that increased es-fMRI brain connectivity is a common trait in people with treatment-resistant focal epilepsy (Pedersen et al., 2015). Similarly, people with Parkinson’s disease undergo a normalisation of functional connectivity similar to healthy controls, during brain stimulation of the subthalamic nucleus (Horn et al., 2019). In combination with our findings, these studies suggest that novel network models may be used to inform the efficacy of brain stimulation in a range of neurological conditions (Halu et al., 2019).

On a group level, we observed decreased network switching during intracranial stimulation also in spatially distant default-mode and frontoparietal networks. There are two plausible explanations of why we observed a reduction in network switching beyond the stimulation sites. The first explanation is based on the inherent nature of group-level research designs. Electrode locations varied between participants in this study (even within the amygdala), and es-fMRI effects from individual stimulation sites may cancel out within a group-level statistical test. As we observed in our exploratory individual-specific analysis, each subject’s network pattern is different with changes in network switching proximate to the electrode location (Figure 6 and Figure A.7). This suggests that es-fMRI group studies in epilepsy can delineate large-scale network effects of brain stimulation whereas individual-level studies may locate brain changes related to distinct stimulation sites. For example, what network properties are affected by specific cortical or sub-cortical stimulation sites in individual subjects, and what is their relationship with individual clinical symptomatology and treatment efficacy? Improving individual-level prediction from quantitative MRI data is needed to provide an answer to these questions. The second explanation is that spatial distortion induced by the intracranial electrodes attenuate the fMRI signal near the stimulation sites (Lee et al., 2012). We are encouraged by recent advances in the field demonstrating that graphene fibre electrodes reduce MRI distortion proximate to the stimulation site (Zhao et al., 2020). Graphene fibre electrodes will benefit future es-fMRI and brain stimulation studies as we seek to further understand local and global brain network variability in response to intracranial brain stimulation.

Another limitation of this study is the relatively short duration of the es-fMRI scans (typically 9 minutes in length). This precluded testing of the long-lasting network effects of intracranial brain stimulation. However, we observed a sudden drop in brain network switching in the early epochs of the es-fMRI scans, followed by a plateau in brain network switching, suggesting that the initial effects of brain stimulation may persist for at least 10 minutes (Figure 3). Longer es-fMRI scans are needed to clarify the duration of stimulation-induced brain network effects A longer period of es-fMRI recording before the onset of the first brain stimulation (here, ~30 seconds into the scan) is also beneficial to ensure the participants are settled and relaxed before the onset of brain stimulation, and it would allow for improved control of filter-related issues that can occur at the start/end of es-fMRI signals.

## Conclusions

Our results suggest that large-scale networks are dynamically modulated by neuronal perturbation induced by intracranial brain stimulation, affecting brain regions and networks distant from the stimulation target. We believe that this research is a necessary first step to enable robust investigation into individual-level network changes following intracranial brain stimulation, as this may aid the clinical decision support of refractory neurological diseases.

## Acknowledgements

We thank the researchers from the University of Iowa comprehensive epilepsy program (Department of Neurosurgery), as well as their collaborators listed in Oya et al. (2017) and Thompson et al. (2020) for acquiring and freely releasing this dataset. The authors declare no competing interests.

## Notes

### Competing Interest Statement

The authors have declared no competing interest.

## References

1. Adcock, J.E., Wise, R.G., Oxbury, J.M., Oxbury, S.M., and Matthews, P.M. (2003). Quantitative fMRI assessment of the differences in lateralization of language-related brain activation in patients with temporal lobe epilepsy. NeuroImage 18, 423–438.

2. Alhourani, A., McDowell, M.M., Randazzo, M.J., Wozny, T.A., Kondylis, E.D., Lipski, W.J., Beck, S., Karp, J.F., Ghuman, A.S., and Richardson, R.M. (2015). Network effects of deep brain stimulation. J. Neurophysiol. 114, 2105–2117.

3. Aksenov, D.P., Li, L., Miller, M.J., and Wyrwicz, A.M. (2019). Role of the inhibitory system in shaping the BOLD fMRI response. NeuroImage 201, 116034.

4. Bassett, D.S., Wymbs, N.F., Porter, M.A., Mucha, P.J., Carlson, J.M., and Grafton, S.T. (2011). Dynamic reconfiguration of human brain networks during learning. Proc. Natl. Acad. Sci. U. S. A. 108, 7641–7646.

5. Bassett, D.S., Porter, M.A., Wymbs, N.F., Grafton, S.T., Carlson, J.M., and Mucha, P.J. (2013). Robust detection of dynamic community structure in networks. Chaos Woodbury N 23, 013142.

6. Bedrosian, E. (1963). A product theorem for Hilbert transforms. Proc. IEEE 51, 868–869.

7. Benazzouz, A., Piallat, B., Pollak, P., and Benabid, A.-L. (1995). Responses of substantia nigra pars reticulata and globus pallidus complex to high frequency stimulation of the subthalamic nucleus in rats: electrophysiological data. Neurosci. Lett. 189, 77–80.

8. Benjamini, Y., and Hochberg, Y. (1995). Controlling the False Discovery Rate: A Practical and Powerful Approach to Multiple Testing. J. R. Stat. Soc. Ser. B Methodol. 57, 289–300.

9. Bernhardt, B.C., Chen, Z., He, Y., Evans, A.C., and Bernasconi, N. (2011). Graph-Theoretical Analysis Reveals Disrupted Small-World Organization of Cortical Thickness Correlation Networks in Temporal Lobe Epilepsy. Cereb. Cortex bhq291.

10. Bernhardt, B.C., Hong, S.-J., Bernasconi, A., and Bernasconi, N. (2015). Magnetic resonance imaging pattern learning in temporal lobe epilepsy: classification and prognostics. Ann. Neurol. 77, 436–446.

11. Betzel, R.F., and Bassett, D.S. (2017). Multi-scale brain networks. NeuroImage 160, 73–83.

12. Betzel, R.F., Satterthwaite, T.D., Gold, J.I., and Bassett, D.S. (2017). Positive affect, surprise, and fatigue are correlates of network flexibility. Sci. Rep. 7, 520.

13. Blahak, C., Bäzner, H., Capelle, H.-H., Wöhrle, J.C., Weigel, R., Hennerici, M.G., and Krauss, J.K. (2009). Rapid response of parkinsonian tremor to STN-DBS changes: Direct modulation of oscillatory basal ganglia activity? Mov. Disord. 24, 1221–1225.

14. Blondel, V.D., Guillaume, J.-L., Lambiotte, R., and Lefebvre, E. (2008). Fast unfolding of communities in large networks. J. Stat. Mech. Theory Exp. 2008, P10008.

15. Braun, U., Schäfer, A., Walter, H., Erk, S., Romanczuk-Seiferth, N., Haddad, L., Schweiger, J.I., Grimm, O., Heinz, A., Tost, H., et al. (2015). Dynamic reconfiguration of frontal brain networks during executive cognition in humans. Proc. Natl. Acad. Sci. U. S. A. 112, 11678–11683.

16. De Domenico, M. (2017). Multilayer modeling and analysis of human brain networks. GigaScience 6, 1–8.

17. Dwivedi, A.K., and Vyas, O.P. (2011). An Exploratory Study of Experimental Tools for Wireless Sensor Networks. Wirel. Sens. Netw. 03, 215–240.

18. Finc, K., Bonna, K., He, X., Lydon-Staley, D.M., Kühn, S., Duch, W., and Bassett, D.S. (2020). Dynamic reconfiguration of functional brain networks during working memory training. Nat. Commun. 11, 2435.

19. Fisher, R.S., Cross, J.H., French, J.A., Higurashi, N., Hirsch, E., Jansen, F.E., Lagae, L., Moshé, S.L., Peltola, J., Perez, E.R., et al. (2017). Operational classification of seizure types by the International League Against Epilepsy: Position Paper of the ILAE Commission for Classification and Terminology. Epilepsia 58, 522–530.

20. Fisher, R., Salanova, V., Witt, T., Worth, R., Henry, T., Gross, R., Oommen, K., Osorio, I., Nazzaro, J., Labar, D., et al. (2010). Electrical stimulation of the anterior nucleus of the thalamus for treatment of refractory epilepsy. Epilepsia 51, 899–908.

21. Fornito, A., Zalesky, A., Pantelis, C., and Bullmore, E.T. (2012). Schizophrenia, neuroimaging and connectomics. NeuroImage 62, 2296–2314.

22. Garcia, L., D’Alessandro, G., Bioulac, B., and Hammond, C. (2005). High-frequency stimulation in Parkinson’s disease: more or less? Trends Neurosci. 28, 209–216.

23. Gerraty, R.T., Davidow, J.Y., Foerde, K., Galvan, A., Bassett, D.S., and Shohamy, D. (2018). Dynamic Flexibility in Striatal-Cortical Circuits Supports Reinforcement Learning. J. Neurosci. 38, 2442–2453.

24. Gifford, G., Crossley, N., Kempton, M.J., Morgan, S., Dazzan, P., Young, J., and McGuire, P. (2020). Resting state fMRI based multilayer network configuration in patients with schizophrenia. NeuroImage Clin. 25, 102169.

25. Glasser, M.F., Coalson, T.S., Robinson, E.C., Hacker, C.D., Harwell, J., Yacoub, E., Ugurbil, K., Andersson, J., Beckmann, C.F., Jenkinson, M., et al. (2016). A multi-modal parcellation of human cerebral cortex. Nature 536, 171–178.

26. Glerean, E., Salmi, J., Lahnakoski, J.M., Jaaskelainen, I.P., and Sams, M. (2012). Functional Magnetic Resonance Imaging Phase Synchronization as a Measure of Dynamic Functional Connectivity. Brain Connect. 2, 91–101.

27. Good, B.H., de Montjoye, Y.-A., and Clauset, A. (2010). Performance of modularity maximization in practical contexts. Phys. Rev. E 81, 046106.

28. Gummadavelli, A., Kundishora, A.J., Willie, J.T., Andrews, J.P., Gerrard, J.L., Spencer, D.D., and Blumenfeld, H. (2015). Neurostimulation to improve level of consciousness in patients with epilepsy. Neurosurg. Focus 38, E10.

29. Halu, A., De Domenico, M., Arenas, A., and Sharma, A. (2019). The multiplex network of human diseases. Npj Syst. Biol. Appl. 5, 1–12.

30. Harlalka, V., Bapi, R.S., Vinod, P.K., and Roy, D. (2019). Atypical Flexibility in Dynamic Functional Connectivity Quantifies the Severity in Autism Spectrum Disorder. Front. Hum. Neurosci. 13, 6.

31. van den Heuvel, M.P., and Sporns, O. (2011). Rich-Club Organization of the Human Connectome. J. Neurosci. 31, 15775–15786.

32. van den Heuvel, M.P., de Lange, S.C., Zalesky, A., Seguin, C., Yeo, B.T.T., and Schmidt, R. (2017). Proportional thresholding in resting-state fMRI functional connectivity networks and consequences for patient-control connectome studies: Issues and recommendations. NeuroImage 152, 437–449.

33. Honari, H., Choe, A.S., and Lindquist, M.A. (2020). Evaluating phase synchronization methods in fMRI: a comparison study and new approaches. NeuroImage, 228, 117704.

34. Hong, S.-J., Bernhardt, B.C., Caldairou, B., Hall, J.A., Guiot, M.C., Schrader, D., Bernasconi, N., and Bernasconi, A. (2017). Multimodal MRI profiling of focal cortical dysplasia type II. Neurology 88, 734–742.

35. Horn, A., Wenzel, G., Irmen, F., Huebl, J., Li, N., Neumann, W.-J., Krause, P., Bohner, G., Scheel, M., and Kühn, A.A. (2019). Deep brain stimulation induced normalization of the human functional connectome in Parkinson’s disease. Brain 142, 3129–3143.

36. Jakobs, M., Fomenko, A., Lozano, A.M., and Kiening, K.L. (2019). Cellular, molecular, and clinical mechanisms of action of deep brain stimulation-a systematic review on established indications and outlook on future developments. EMBO Mol. Med. 11.

37. Johnson, M.D., Miocinovic, S., McIntyre, C.C., and Vitek, J.L. (2008). Mechanisms and targets of deep brain stimulation in movement disorders. Neurotherapeutics 5, 294–308.

38. Khan, S., Wright, I., Javed, S., Sharples, P., Jardine, P., Carter, M., and Gill, S.S. (2009). High frequency stimulation of the mamillothalamic tract for the treatment of resistant seizures associated with hypothalamic hamartoma. Epilepsia 50, 1608–1611.

39. Lafreniere-Roula, M., Kim, E., Hutchison, W.D., Lozano, A.M., Hodaie, M., and Dostrovsky, J.O. (2010). High-frequency microstimulation in human globus pallidus and substantia nigra. Exp. Brain Res. 205, 251–261.

40. Lancichinetti, A., and Fortunato, S. (2012). Consensus clustering in complex networks. Sci. Rep. 2, 336.

41. Langer, N., Pedroni, A., and Jäncke, L. (2013). The Problem of Thresholding in Small-World Network Analysis. PLoS ONE 8, e53199.

42. Lee, K.J., Shon, Y.M., and Cho, C.B. (2012). Long-term outcome of anterior thalamic nucleus stimulation for intractable epilepsy. Stereotact. Funct. Neurosurg. 90, 379–385.

43. Li, D.-H., and Yang, X.-F. (2017). Remote modulation of network excitability during deep brain stimulation for epilepsy. Seizure 47, 42–50.

44. Li, M.C.H., and Cook, M.J. (2018). Deep brain stimulation for drug-resistant epilepsy. Epilepsia 59, 273–290.

45. Li, Q., Wang, X., Wang, S., Xie, Y., Li, X., Xie, Y., and Li, S. (2019). Dynamic reconfiguration of the functional brain network after musical training in young adults. Brain Struct. Funct. 224, 1781–1795.

46. Long, Y., Chen, C., Deng, M., Huang, X., Tan, W., Zhang, L., Fan, Z., and Liu, Z. (2019). Psychological resilience negatively correlates with resting-state brain network flexibility in young healthy adults: a dynamic functional magnetic resonance imaging study. Ann. Transl. Med. 7.

47. Lydon-Staley, D.M., Ciric, R., Satterthwaite, T.D., and Bassett, D.S. (2018). Evaluation of confound regression strategies for the mitigation of micromovement artifact in studies of dynamic resting-state functional connectivity and multilayer network modularity. Netw. Neurosci. 3, 427–454.

48. Middlebrooks, E.H., Grewal, S.S., Stead, M., Lundstrom, B.N., Worrell, G.A., and Van Gompel, J.J. (2018). Differences in functional connectivity profiles as a predictor of response to anterior thalamic nucleus deep brain stimulation for epilepsy: a hypothesis for the mechanism of action and a potential biomarker for outcomes. Neurosurg. Focus 45, E7.

49. Mormann, F., Lehnertz, K., David, P., and E. Elger, C. (2000). Mean phase coherence as a measure for phase synchronization and its application to the EEG of epilepsy patients. Phys. Nonlinear Phenom. 144, 358–369.

50. Morrell, M.J., and RNS System in Epilepsy Study Group (2011). Responsive cortical stimulation for the treatment of medically intractable partial epilepsy. Neurology 77, 1295–1304.

51. Mucha, P.J., Richardson, T., Macon, K., Porter, M.A., and Onnela, J.-P. (2010). Community structure in time-dependent, multiscale, and multiplex networks. Science 328, 876–878.

52. Oya, H., Howard, M.A., Magnotta, V.A., Kruger, A., Griffiths, T.D., Lemieux, L., Carmichael, D.W., Petkov, C.I., Kawasaki, H., Kovach, C.K., et al. (2017). Mapping effective connectivity in the human brain with concurrent intracranial electrical stimulation and BOLD-fMRI. J. Neurosci. Methods 277, 101–112.

53. Paban, V., Modolo, J., Mheich, A., and Hassan, M. (2019). Psychological resilience correlates with EEG source-space brain network flexibility. Netw. Neurosci. 3, 539–550.

54. Pedersen, M., Omidvarnia, A.H., Walz, J.M., and Jackson, G.D. (2015). Increased segregation of brain networks in focal epilepsy: An fMRI graph theory finding. NeuroImage Clin. 8, 536–542.

55. Pedersen, M., Curwood, E.K., Vaughan, D.N., Omidvarnia, A.H., and Jackson, G.D. (2016). Abnormal Brain Areas Common to the Focal Epilepsies: Multivariate Pattern Analysis of fMRI. Brain Connect. 6, 208–215.

56. Pedersen, M., Omidvarnia, A., Curwood, E.K., Walz, J.M., Rayner, G., and Jackson, G.D. (2017). The dynamics of functional connectivity in neocortical focal epilepsy. NeuroImage Clin. 15, 209–214.

57. Pedersen, M., Zalesky, A., Omidvarnia, A., and Jackson, G.D. (2018a). Multilayer network switching rate predicts brain performance. Proc. Natl. Acad. Sci. 115, 13376–13381.

58. Pedersen, M., Omidvarnia, A., Zalesky, A., and Jackson, G.D. (2018b). On the relationship between instantaneous phase synchrony and correlation-based sliding windows for time-resolved fMRI connectivity analysis. NeuroImage 181, 85–94.

59. Ponce-Alvarez, A., Deco, G., Hagmann, P., Romani, G.L., Mantini, D., and Corbetta, M. (2015). Resting-State Temporal Synchronization Networks Emerge from Connectivity Topology and Heterogeneity. PLoS Comput Biol 11, e1004100.

60. Sarzynska, M., Leicht, E.A., Chowell, G., and Porter, M.A. (2016). Null models for community detection in spatially embedded, temporal networks. J. Complex Netw. 4, 363–406.

61. Schulze-Bonhage, A. (2017). Brain stimulation as a neuromodulatory epilepsy therapy. Seizure 44, 169–175.

62. Shao, J., Dai, Z., Zhu, R., Wang, X., Tao, S., Bi, K., Tian, S., Wang, H., Sun, Y., Yao, Z., et al. (2019). Early identification of bipolar from unipolar depression before manic episode: Evidence from dynamic rfMRI. Bipolar Disord. 21, 774–784.

63. Shine, J.M., Koyejo, O., and Poldrack, R.A. (2016). Temporal metastates are associated with differential patterns of time-resolved connectivity, network topology, and attention. Proc. Natl. Acad. Sci. 113, 9888–9891.

64. Sporns, O., and Betzel, R.F. (2016). Modular Brain Networks. Annu. Rev. Psychol. 67, 613–640.

65. Telesford, Q.K., Ashourvan, A., Wymbs, N.F., Grafton, S.T., Vettel, J.M., and Bassett, D.S. (2017). Cohesive network reconfiguration accompanies extended training. Hum. Brain Mapp. 38, 4744–4759.

66. Thompson, W.H., Esteban, O., Oya, H., Nair, R., Eberhardt, F., Dubois, J., Poldrack, R.A., Adolphs, R., and Shine, J.M. (2021). Intracranial electrical stimulation alters meso-scale network integration as a function of network topology. BioRxiv 2021.01.16.426941.

67. Thompson, W., Nair, R., Oya, H., Esteban, O., Shine, J., Petkov, C., Poldrack, R., Howard, M., and Adolphs, R. (2020). Human es-fMRI Resource: Concurrent deep-brain stimulation and whole-brain functional MRI. bioRxiv. doi: https://doi.org/10.1101/2020.05.18.102657.

68. Tian, S., Sun, Y., Shao, J., Zhang, S., Mo, Z., Liu, X., Wang, Q., Wang, L., Zhao, P., Chattun, M.R., et al. (2020a). Predicting escitalopram monotherapy response in depression: The role of anterior cingulate cortex. Hum. Brain Mapp. 41, 1249–1260.

69. Tian, Y., Margulies, D.S., Breakspear, M., and Zalesky, A. (2020b). Topographic organization of the human subcortex unveiled with functional connectivity gradients. Nat. Neurosci. 23, 1421–1432.

70. Toprani, S., and Durand, D.M. (2013). Long-lasting hyperpolarization underlies seizure reduction by low frequency deep brain electrical stimulation. J. Physiol. 591, 5765–5790.

71. Vaiana, M., and Muldoon, S. (2018). Multilayer Brain Networks. J. Nonlinear Sci.

72. Yang, Z., Telesford, Q.K., Franco, A.R., Lim, R., Gu, S., Xu, T., Ai, L., Castellanos, F.X., Yan, C.-G., Colcombe, S., et al. (2020). Measurement Reliability for Individual Differences in Multilayer Network Dynamics: Cautions and Considerations. NeuroImage 117489.

73. Yeo, B.T.T., Krienen, F.M., Sepulcre, J., Sabuncu, M.R., Lashkari, D., Hollinshead, M., Roffman, J.L., Smoller, J.W., Zöllei, L., Polimeni, J.R., et al. (2011). The organization of the human cerebral cortex estimated by intrinsic functional connectivity. J. Neurophysiol. 106, 1125–1165.

74. Zangiabadi, N., Ladino, L.D., Sina, F., Orozco-Hernández, J.P., Carter, A., and Téllez-Zenteno, J.F. (2019). Deep Brain Stimulation and Drug-Resistant Epilepsy: A Review of the Literature. Front. Neurol. 10.

75. Zhao, S., Li, G., Tong, C., Chen, W., Wang, P., Dai, J., Fu, X., Xu, Z., Liu, X., Lu, L., et al. (2020). Full activation pattern mapping by simultaneous deep brain stimulation and fMRI with graphene fiber electrodes. Nat. Commun. 11, 1788.

