## Supplementary material for "Intracranial brain stimulation modulates fMRI-based network switching": Figure A.

**Appendix A: fMRI pre-processing:**

Pre-processing performed using fMRIPrep1.5.1rc1 (RRID: SCR_016216 - Esteban et al., 2019, 2020), which is based on Nipype 1.3.0-rc1 (RRID: SCR_002502 - Gorgolewski et al., 2011).

***Anatomical MRI pre-processing***

The T1-weighted (T1w) image was corrected for intensity non-uniformity (INU) with N4BiasFieldCorrection (Tustison et al., 2010), distributed with ANTs 2.2.0 (RRID: SCR_004757) (Avants et al., 2008), and used as T1w-reference throughout the workflow. The T1w-reference was then skull-stripped with a Nipype implementation of the antsBrainExtraction.sh workflow (from ANTs), using OASIS30ANTs as a target template. Brain tissue segmentation of cerebrospinal fluid (CSF), white matter (WM) and grey matter (GM) were performed on the brain extracted T1w using fast (FSL 5.0.9, RRID: SCR_002823 - Zhang et al., 2001). Brain surfaces were reconstructed using recon-all (FreeSurfer 6.0.1, RRID: SCR_001847 - Dale et al., 1999), and the brain mask estimated previously was refined with a custom variation of the method to reconcile ANTs-derived and FreeSurfer-derived segmentation of the cortical grey-matter of Mindboggle (RRID: SCR_002438 - Klein et al., 2017). Volume-based spatial normalization to one standard space (MNI152NLin2009cAsym) was performed through nonlinear registration with antsRegistration (ANTs 2.2.0), using brain-extracted versions of both T1w reference and the T1w template. The following template was selected for spatial normalization: ICBM 152 Nonlinear Asymmetrical template version 2009c (RRID:SCR_008796; TemplateFlow ID: [MNI152NLin2009cAsym] - Fonov et al., 2009).

***es-fMRI pre-processing***

For each of the BOLD runs found per subject (across all tasks and sessions), the following pre-processing was performed. First, a reference volume and its skull-stripped version were generated using a custom methodology of fMRIPrep. A deformation field to correct for susceptibility distortions was estimated based on fMRIPrep's field map-less approach. The deformation field is that resulting from co-registering the BOLD reference to the same-subject T1w-reference with its intensity inverted (Huntenburg, 2014; Wang et al., 2017). Registration was performed with antsRegistration (ANTs 2.2.0), and the process regularized by constraining deformation to be nonzero only along the phase-encoding direction and modulated with an average field map template (Treiber et al., 2016). Based on the estimated susceptibility distortion, an unwarped BOLD reference was calculated for a more accurate co-registration with the anatomical reference. The BOLD reference was then co-registered to the T1w reference using bbregister (FreeSurfer) which implements boundary-based registration (Greve and Fischl, 2009). Co-registration was configured with six degrees of freedom.

Head-motion parameters for the BOLD reference (transformation matrices, and six corresponding rotation and translation parameters) were estimated before any spatiotemporal filtering using mcflirt (FSL 5.0.9) (Jenkinson et al., 2002). BOLD runs were slice-time corrected using 3dTshift from AFNI 20160207 (RRID: SCR_005927 - Cox and Hyde, 1997). The BOLD time-series were resampled to surfaces on the following spaces: fsnative, fsaverage5. The BOLD time-series (including slice-timing correction when applied) were resampled onto their original, native space by applying a single, composite transform to correct for head-motion and susceptibility distortions. These resampled BOLD time-series will be referred to as pre-processed BOLD in original space, or just pre-processed BOLD. The BOLD time-series were resampled into standard space, generating a pre-processed BOLD run in [‘MNI152NLin2009cAsym’] space. First, a reference volume and its skull-stripped version were generated using a custom methodology of fMRIPrep. Several confounding time-series were calculated based on the pre-processed BOLD: framewise displacement (FD), DVARS and three region-wise global signals. FD and DVARS are calculated for each functional run, both using their implementations in Nipype (following the definitions in Power et al., 2014). The three global signals are extracted within the CSF, the WM, and the whole-brain masks. Additionally, a set of physiological regressors were extracted to allow for component-based noise correction (CompCor) (Behzadi et al., 2007). Principal components are estimated after high-pass filtering the pre-processed BOLD time-series (using a discrete cosine filter with 128-second cut-off) for the two CompCor variants: temporal (tCompCor) and anatomical (aCompCor). tCompCor components are then calculated from the top 5% variable voxels within a mask covering the subcortical regions. This subcortical mask is obtained by heavily eroding the brain mask, which ensures it does not include cortical GM regions.

For aCompCor, components are calculated within the intersection of the aforementioned mask and the union of CSF and WM masks calculated in T1w space, after their projection to the native space of each functional run (using the inverse BOLD-to-T1w transformation). Components are also calculated separately within the WM and CSF masks. For each CompCor decomposition, the k components with the largest singular values are retained, such that the retained components’ time-series are sufficient to explain 50 percent of variance across the nuisance mask (CSF, WM, combined, or temporal). The remaining components are dropped from consideration. The head-motion estimates calculated in the correction step were also placed within the corresponding confounds file. The confound time series derived from head motion estimates and global signals were expanded with the inclusion of temporal derivatives and quadratic terms for each (Satterthwaite et al., 2013). Frames that exceeded a threshold of 0.5 mm FD or 1.5 standardised DVARS were annotated as motion outliers. All resamplings can be performed with a single interpolation step by composing all the pertinent transformations (i.e. head-motion transform matrices, susceptibility distortion correction, and co-registrations to anatomical and output spaces). Gridded (volumetric) resamplings were performed using antsApplyTransforms (ANTs), configured with Lanczos interpolation to minimize the smoothing effects of other kernels (Lanczos, 1964).

Many internal operations of fMRIPrep use Nilearn 0.5.2 (RRID: SCR_001362 - Abraham et al., 2014), mostly within the functional processing workflow. For more details of the pipeline, see the section corresponding to workflows in fMRIPrep's documentation (https://fmriprep.org//downloads/en/0.2.0/pdf/).

**References:**

Abraham, A., Pedregosa, F., Eickenberg, M., Gervais, P., Mueller, A., Kossaifi, J., Gramfort, A., Thirion, B., and Varoquaux, G. (2014). Machine learning for neuroimaging with scikit-learn. Front. Neuroinformatics *8*.

Avants, B.B., Epstein, C.L., Grossman, M., and Gee, J.C. (2008). Symmetric diffeomorphic image registration with cross-correlation: Evaluating automated labeling of elderly and neurodegenerative brain. Med. Image Anal. *12*, 26–41.

Behzadi, Y., Restom, K., Liau, J., and Liu, T.T. (2007). A component based noise correction method (CompCor) for BOLD and perfusion based fMRI. NeuroImage *37*, 90–101.

Cox, R.W., and Hyde, J.S. (1997). Software tools for analysis and visualization of fMRI data. NMR Biomed. *10*, 171–178.

Dale, A.M., Fischl, B., and Sereno, M.I. (1999). Cortical Surface-Based Analysis: I. Segmentation and Surface Reconstruction. NeuroImage *9*, 179–194.

Esteban, O., Markiewicz, C.J., Blair, R.W., Moodie, C.A., Isik, A.I., Erramuzpe, A., Kent, J.D., Goncalves, M., DuPre, E., Snyder, M., et al. (2019). fMRIPrep: a robust preprocessing pipeline for functional MRI. Nat. Methods *16*, 111–116.

Esteban, O., Markiewicz, C.J., Goncalves, M., DuPre, E., Kent, J.D., Salo, T., Ciric, R., Pinsard, B., Blair, R.W., Poldrack, R.A., et al. (2020). fMRIPrep: a robust preprocessing pipeline for functional MRI (Zenodo).

Fonov, V., Evans, A., McKinstry, R., Almli, C., and Collins, D. (2009). Unbiased nonlinear average age-appropriate brain templates from birth to adulthood. NeuroImage *47*, S102.

Gorgolewski, K., Burns, C.D., Madison, C., Clark, D., Halchenko, Y.O., Waskom, M.L., and Ghosh, S.S. (2011). Nipype: A Flexible, Lightweight and Extensible Neuroimaging Data Processing Framework in Python. Front. Neuroinformatics *5*.

Greve, D.N., and Fischl, B. (2009). Accurate and robust brain image alignment using boundary-based registration. NeuroImage *48*, 63–72.

Huntenburg, J.M. (2014). Evaluating nonlinear coregistration of BOLD EPI and T1w images. Freie Universität Berlin.

Jenkinson, M., Bannister, P., Brady, M., and Smith, S. (2002). Improved Optimization for the Robust and Accurate Linear Registration and Motion Correction of Brain Images. NeuroImage *17*, 825–841.

Klein, A., Ghosh, S.S., Bao, F.S., Giard, J., Häme, Y., Stavsky, E., Lee, N., Rossa, B., Reuter, M., Neto, E.C., et al. (2017). Mindboggling morphometry of human brains. PLOS Comput. Biol. *13*, e1005350

Lanczos, C. (1964). Evaluation of Noisy Data. J. Soc. Ind. Appl. Math. Ser. B Numer. Anal. *1*, 76–85.

Power, J.D., Mitra, A., Laumann, T.O., Snyder, A.Z., Schlaggar, B.L., and Petersen, S.E. (2014). Methods to detect, characterize, and remove motion artifact in resting state fMRI. NeuroImage *84*, 320–341.

Satterthwaite, T.D., Elliott, M.A., Gerraty, R.T., Ruparel, K., Loughead, J., Calkins, M.E., Eickhoff, S.B., Hakonarson, H., Gur, R.C., Gur, R.E., et al. (2013). An improved framework for confound regression and filtering for control of motion artifact in the preprocessing of resting-state functional connectivity data. NeuroImage *64*, 240–256.

Treiber, J.M., White, N.S., Steed, T.C., Bartsch, H., Holland, D., Farid, N., McDonald, C.R., Carter, B.S., Dale, A.M., and Chen, C.C. (2016). Characterization and Correction of Geometric Distortions in 814 Diffusion Weighted Images. PLOS ONE *11*, e0152472.

Tustison, N.J., Avants, B.B., Cook, P.A., Zheng, Y., Egan, A., Yushkevich, P.A., and Gee, J.C. (2010). N4ITK: Improved N3 Bias Correction. IEEE Trans. Med. Imaging *29*, 1310–1320.

Wang, S., Peterson, D.J., Gatenby, J.C., Li, W., Grabowski, T.J., and Madhyastha, T.M. (2017). Evaluation of Field Map and Nonlinear Registration Methods for Correction of Susceptibility Artifacts in Diffusion MRI. Front. Neuroinformatics *11*.

Zhang, Y., Brady, M., and Smith, S. (2001). Segmentation of brain MR images through a hidden Markov random field model and the expectation-maximization algorithm. IEEE Trans. Med. Imaging *20*, 45–57.

### *Figure A.1*

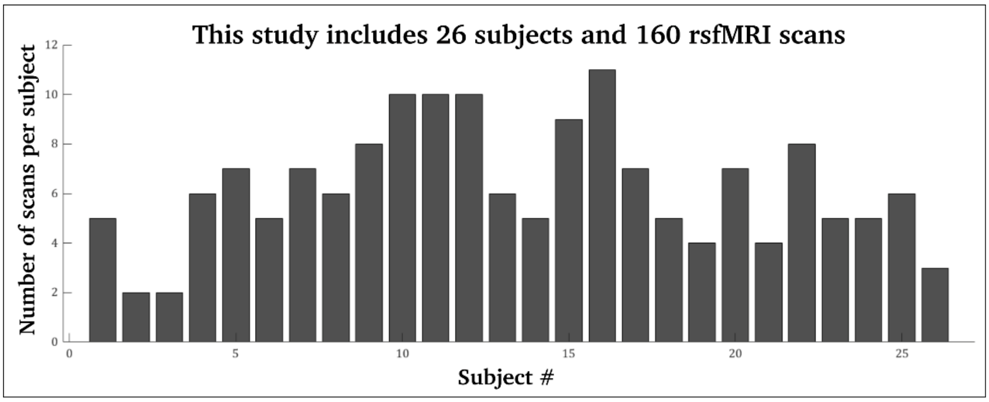


***A1: Number of individuals included in the study (x-axis), and how many fMRI scans were acquired in each subject (y-axis):*** We analysed a total of 160 rsfMRI scans from 26 individuals, and we included rsfMRI scans that were at least 9 minutes long. Multiple rsfMRI scans per subject ensure that results are consistent across different acquisition times and brain states.

***Figure A.2***

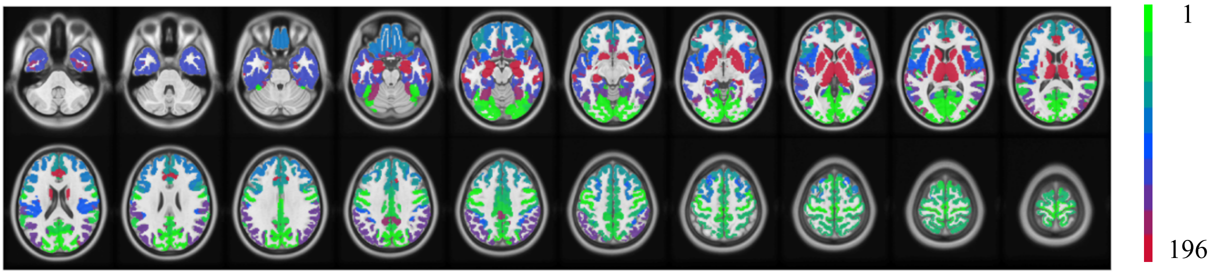


***A2: Brain parcellation mask with symmetric cortical and sub-cortical brain regions:*** Anatomical location of brain regions in the parcellation mask we used in this study. This parcellation mask comprised 196 symmetric cortical and subcortical brain regions, based on a parcellation mask from the Human Connectome Project with 180 cortical brain regions (Glasser et al. 2016). We combined this cortical mask with 16 homogenous subcortical brain regions (Tian et al. 2020).

***Figure A.3***

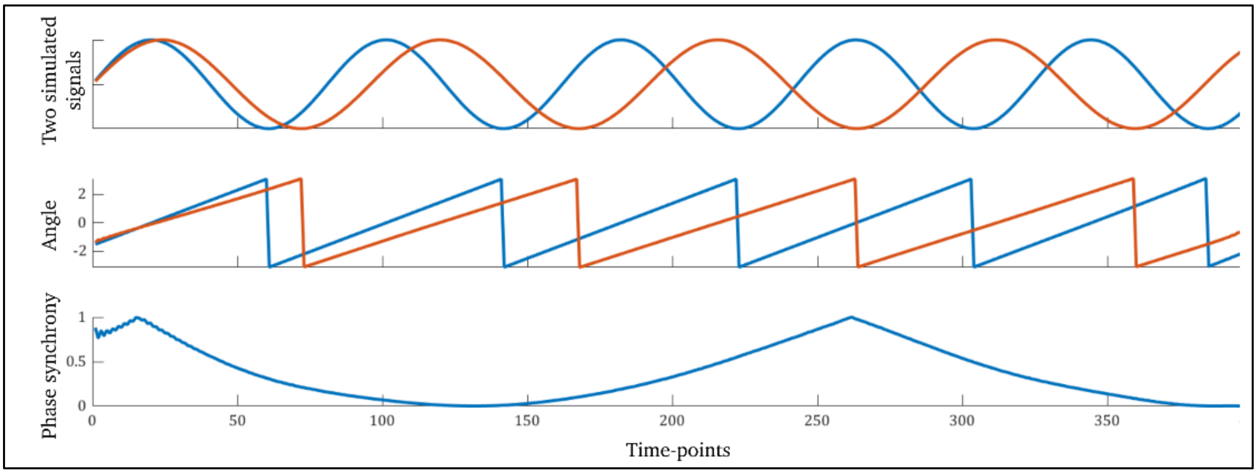


***A3: An example of Instantaneous Phase Synchrony:*** *First row:* We simulated two signals that fluctuated in-and-out of synchrony. *Second row:* Instantaneous phase synchrony is quantified by calculating the angle between signals, using the Hilbert transform. *Third row:* Instantaneous phase synchrony is then entered in the multilayer modularity model.

*Figure A.4*

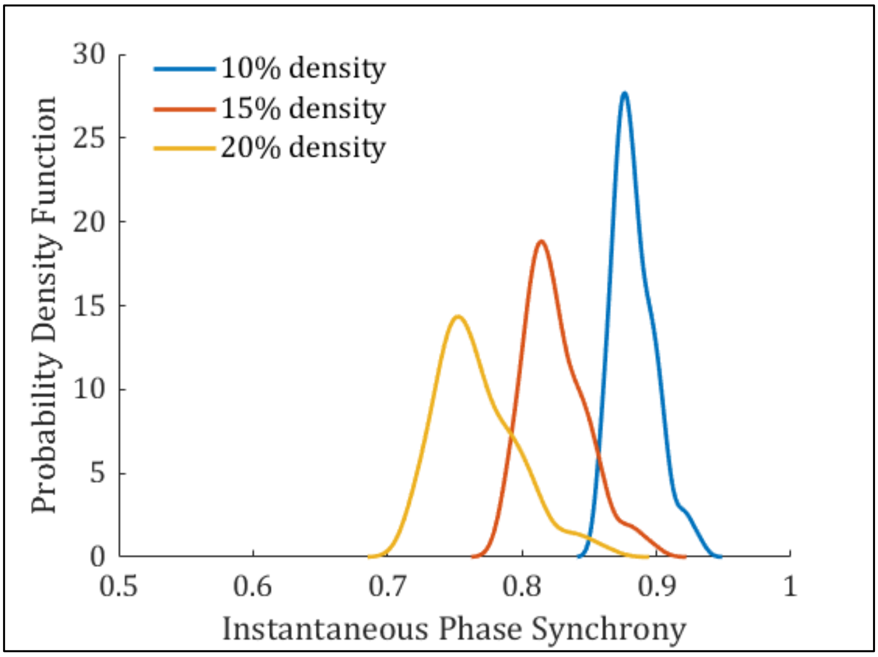


***A4: Minimum phase synchrony values for network thresholding, across 160 rsfMRI scans:*** Displayed are 3 density distributions based on network density thresholds at 10% (blue), 15% (red) and 20% (yellow). The distribution of values corresponds to the minimum instantaneous phase synchrony values included in thresholded spatiotemporal matrices. As reported in the main manuscripts, our findings suggest that lower phase synchrony values (e.g., at a 20% network density) may include more noise and artefacts, and therefore attenuate final statistical results.

***Figure A.5***

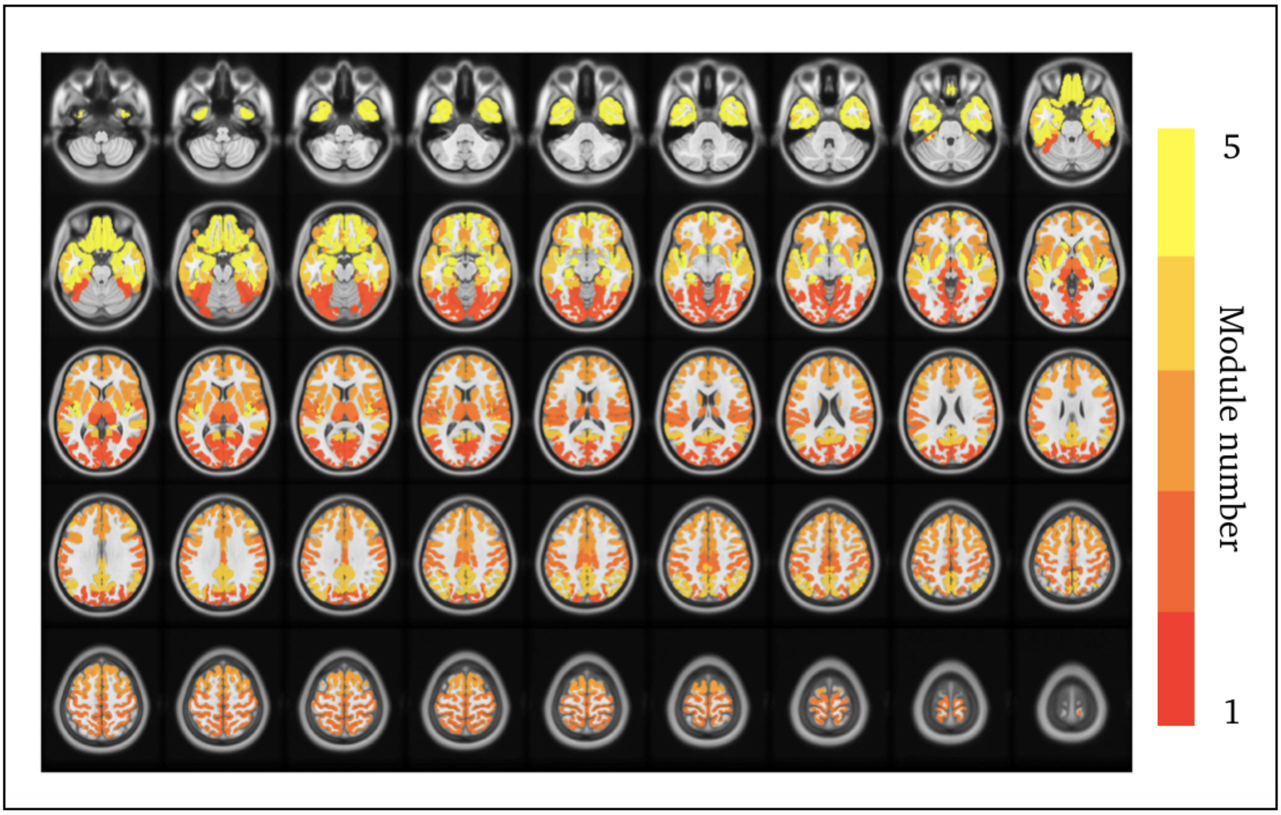


***A5: Consensus clustering across subjects from the multilayer modularity model:*** Here we display the consensus clustering results based on the multilayer modularity models. The consensus algorithm quantifies brain regions that consistently belong to the same modules, across all rsfMRI scans. Five modules were found to have strong intramodular connectivity across all datasets, including, i) visual network, ii) somatomotor network, iii) frontoparietal network, iv) default-mode network and v) sub-cortical brain regions.

*Figure A.6*

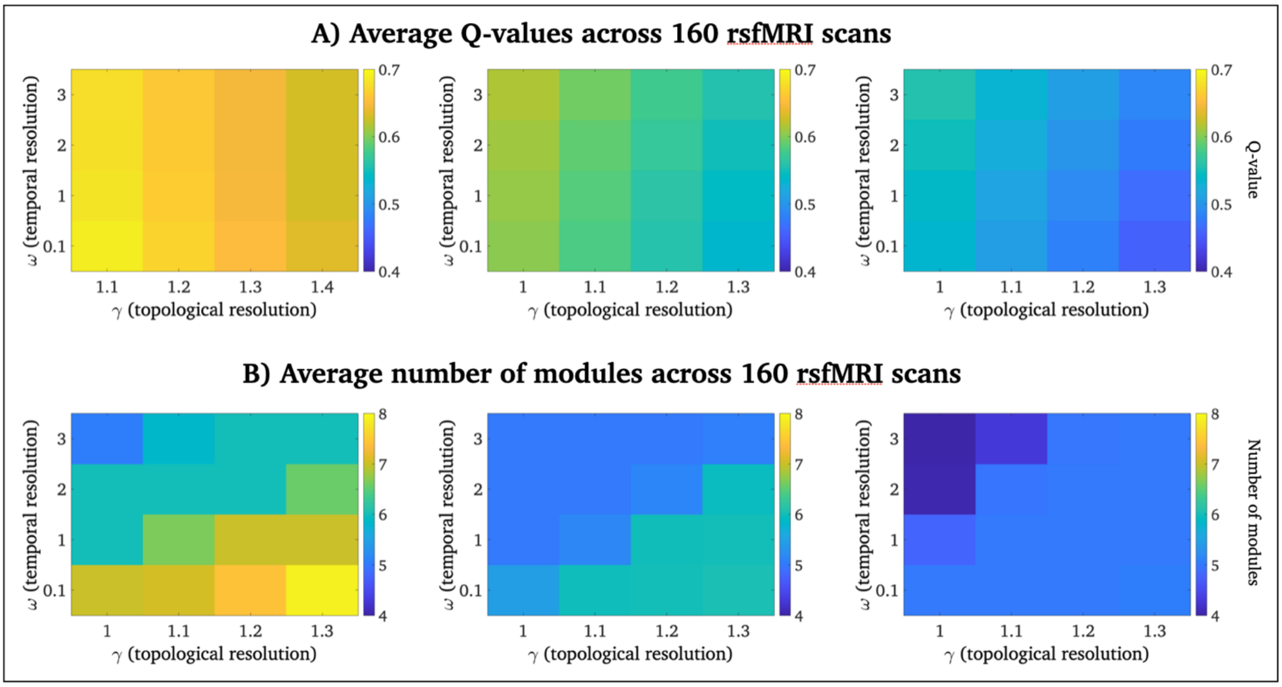


***A6: Average modularity across network densities and network resolution parameters:*** A) This is a representation of average Q-values across all rsfMRI scans. Higher Q-values means that a network is more modular. For example, a Q-value of 1 indicates that nodes within a specific module only have intra-modular connections. Here, low-density networks (10% density), as well as low topological resolution (x-axes) and high temporal resolution (y-axis), are associated with higher modularity. Q-values for 10% network density = 0.68 ± 0.02 s.d.; 15% network density = 0.59 ± 0.02 s.d.; and 20% network density = 0.52 ± 0.03 s.d., across all rsfMRI scans. B) The average number of modules across all rsfMRI scans. We observed that low-density networks (10% density) and higher $\gamma$-values (x-axes) returned more network modules. The average number of modules ranged between 4 and 8.

*Figure A.7*

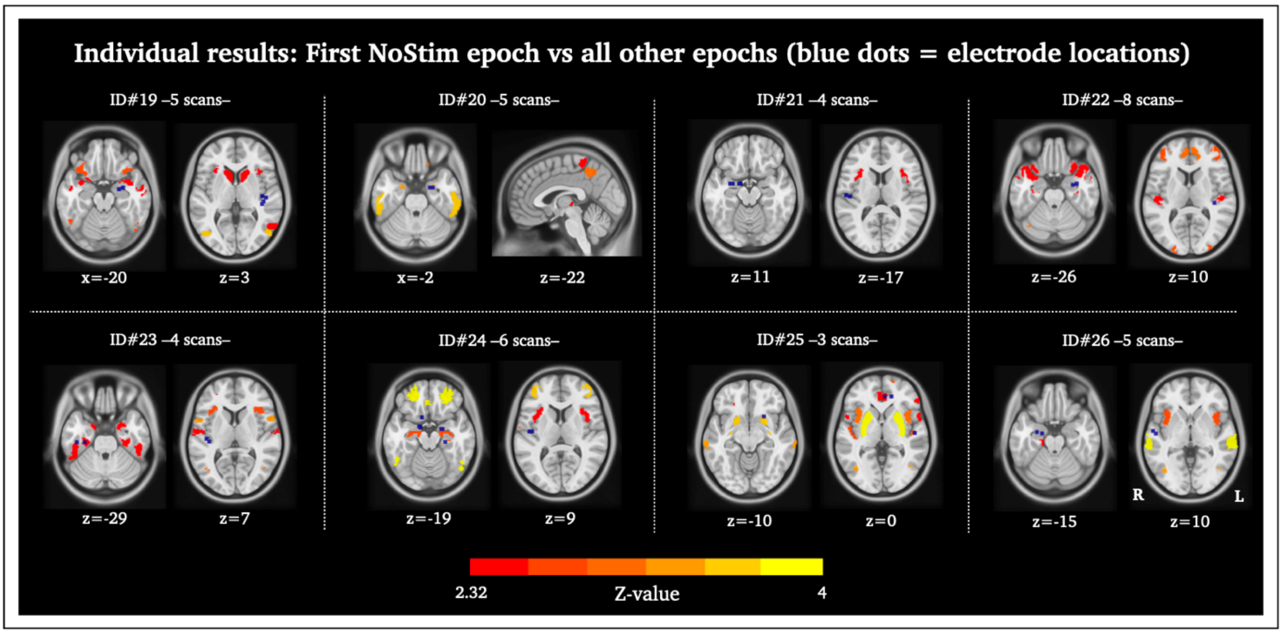


***A7: Individual results – first NoStim epoch versus all other epochs:*** *Here we have presented individual-level decreases in network switching occurring after the first NoStim epoch. Included are the remaining 8 of the 23 epilepsy subjects not presented in Figure 6 in the main manuscript. Blue dots = electrode information.*

### *Figure A.8*

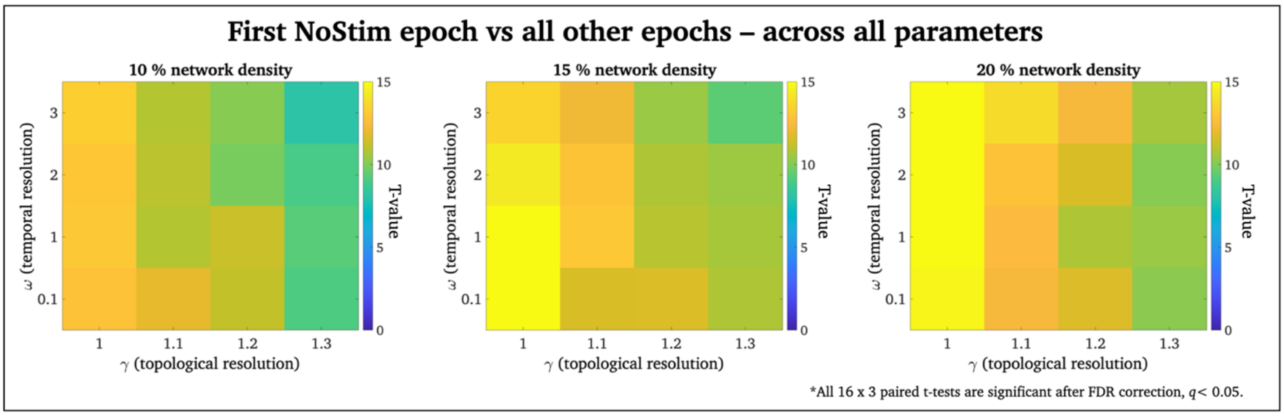


***A8: Effects of spatiotemporal modularity parameters and network density***: Here, we assessed the effects of topological resolution ($\gamma$ – *constraining the size of networks*) and temporal resolution ($\omega$ – *constraining the connectivity strength between adjacent time-points*) in multilayer networks. These parameters are important when interpreting multilayer modularity models and need to be critically assessed while establishing guidelines for optimal multilayer modularity parameters. We replicated our result of reduced network switching during Stim versus NoStim epochs (FDR corrected) for 16 $\gamma/\omega$ parameter combinations ($\gamma$ = 1, 1.1, 1.2, 1.3 and $\omega$ = 0.9, 0.1, 1 We also replicated our analysis for several network density thresholds with a proportional threshold at 10%, 15% (used in our main analyses), and 20% network sparsity. Network density refers to the proportion of connection-pairs that are preserved in the instantaneous phase synchrony networks used as an input to the multilayer modularity model. Post-thresholded matrices were binarized. Although all three network densities were statistically significant between Stim and NoStim epochs (FDR corrected) with a trend towards greater statistical power between Stim and NoStim epochs at higher network density thresholds.

*Figure A.9*

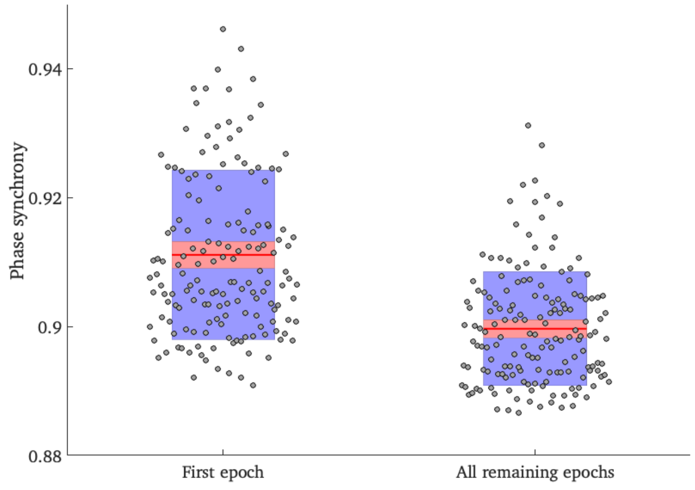


***A9: Reduced phase synchrony during brain stimulation**:* Mean phase synchrony was reduced during Stim epochs compared to NoStim epochs (paired t-test: $t(159) = 10.7$, $p < 0.001$). Each black dot in this figure represents a single rsfMRI scan. The solid red line represents the mean of all scans. The shaded red area is the 95$^{th}$ confidence interval of the mean and the shaded blue area is one standard deviation of the mean.
